# Cracking Controls ATP Hydrolysis in the catalytic unit of a P-type ATPase

**DOI:** 10.1101/2025.01.29.635504

**Authors:** M. Agueda Placenti, Santiago A. Martinez-Gache, Rodolfo M. González-Lebrero, Peter G. Wolynes, F. Luis González Flecha, Ernesto A. Roman

## Abstract

Membrane transporters are essential for homeostasis and among them P-type ATPases are key players. Despite extensive research, conformational changes in their catalytic unit and their coupling to ATP hydrolysis are not explored in detail. In this work, we analyzed the effect of ATP, temperature, and urea on the steady-state ATPase activity, tryptophan fluorescence and far-UV ellipticity of the catalytic unit of the thermophilic Cu(I) transport P_1B_-ATPase from *Archaeoglobus fulgidus*. Combining local frustration analysis with AlphaFold2, we identified an open conformation which we used to perform structure-based model simulations of the open-closed transition. We developed a mechanistic model that fully describes all of our experimental observations. Our results revealed a “cracking”-like mechanism involved in the catalysis of ATP hydrolysis. These findings reinforce that, although simple, the isolated catalytic unit is a relevant model to study the role of local unfolding in the catalytic mechanism of these proteins.

## Introduction

P-type ATPases constitute a large and ubiquitous family of membrane proteins and, based on their substrate specificity and sequence similarity, are classified in P_1_-P_5_ subfamilies ^1^. They are responsible for the active transport of substrates across biological membranes by coupling the highly exergonic ATP hydrolysis to a series of conformational changes resulting in substrate transport ^1–4^. Most of the structural and mechanistic information comes from two P_2_-ATPases: the Ca^2+^-ATPase from sarcoplasmic reticulum (SERCA) ^5–10^ and the Na^+^,K^+^-ATPase ^10–13^. In addition, the Cu(I)-ATPases, which belong to the P_1B_ subfamily involved in heavy metal ion homeostasis, have been the subject of a significant amount of studies ^14–17^. The general reaction mechanism for all P-ATPases is described by the Post-Albers cycle (Figure S1), and includes ATP binding, hydrolysis and the formation of a phosphorylated intermediate ^2^. This cycle involves two main ensembles of conformations named E1 and E2, a definition based on their proteolytic patterns, with the ligand binding site facing either the inward or outward side of the membrane^18^.

Although each subfamily has its own structural characteristics, all members share a common topology composed by a transmembrane domain, through which the substrate is transported, and a series of cytoplasmic domains termed as A (catalytic modulator or “actuator”), N (“nucleotide binding”) and P (“phosphorylation”) ^1,19^ (Figure S2). In particular, P_1B_-ATPases also present a variable number of metal ion binding cytosolic domains (MBDs) ^20^. The conserved N and P domains comprise a catalytic unit (NP) which has been studied in an isolated form, both structurally and functionally, for a small number of P-type ATPases ^21–28^. The observed structures of the isolated NP unit from Cu(I) transport ATPases have been classified as closed when the orientation of helices 5 and 7 is quasi parallel, and open when these helices adopt a perpendicular orientation ^29^. Analogously, in the case of the SERCA, open and closed conformations of its catalytic unit are defined by the orientation of helices 7 and 10 (e.g. PDB ID 1SU4 and 4XOU, respectively) ^30,31^. The existence of these alternative states is associated with the exposure of the ATP-binding cleft, which facilitates ATP binding, and the subsequent closure of the catalytic unit to enable the hydrolysis reaction. In the context of the full-length protein, the closure of the catalytic unit favors the interaction between the A and N domains. This process involves precise coordination between phosphorylation, A domain rotation and transmembrane domain rearrangements mediated through critical interdomain interactions. In P_1B_-ATPases, their unique structural features introduce additional regulatory complexity in heavy metal transport ^10,20^. For example, in the case of the Cu(I) ATPases, MBDs have been recently demonstrated to play a key role in transport regulation depending on the intracellular copper concentration ^20^. Even though the coupling between conformational changes and transport has been largely studied, the complexity of this mechanism is not fully understood ^4,10,19,33^. Thus, studying how the large rearrangement of the N and P domains is coupled to ATP hydrolysis becomes relevant, and the catalytic unit offers a simple yet specific model for this purpose.

Large rearrangements in proteins are usually related to highly dynamic regions ^34^. According to the “principle of minimal frustration”, proteins minimize energetic conflicts in their structures so that they can fold properly while keeping functional conformational dynamics ^35^. Usually, these energetic conflicts are localized in regions involved in conformational changes. In many cases, the limiting step for the rearrangement is a local unfolding event, through a process commonly named as “cracking” which enables structural flexibility and functional motions to happen ^36–40^. A way to experimentally test this mechanism has been the use of chaotropes to lower the activation barrier for the local unfolding step and consequently accelerate the catalyzed reaction ^37,41^. Recently, a novel approach combining local frustration analysis and structure prediction using AlphaFold2 succeeded in describing the conformational transitions of adenylate kinase linked to phosphate transfer between AMP and ATP molecules ^42^.

In this work, we studied the open-closed transition with the aid of experimental and computational tools highlighting the role of strain and “cracking” in the enzyme catalysis. Our results suggest a role of cracking in temperature optimum activity regulation.

## Results and Discussion

### Temperature differentially affects the ATP apparent affinity and the catalytic constant of the enzyme-catalyzed ATP hydrolysis

As was mentioned, *Af*CopA-NP is a catalytic unit formed by two of the soluble domains of the Cu(I) transport P-type ATPase from the hyperthermophilic organism *Archaeoglobus fulgidus*, *Af*CopA. To assess whether this unit, when isolated, conserves its native features and to gain insights into its catalytic mechanism, we evaluated the effect of temperature and substrate concentration on the steady state ATP hydrolysis (Figure 1).

**Figure 1.**
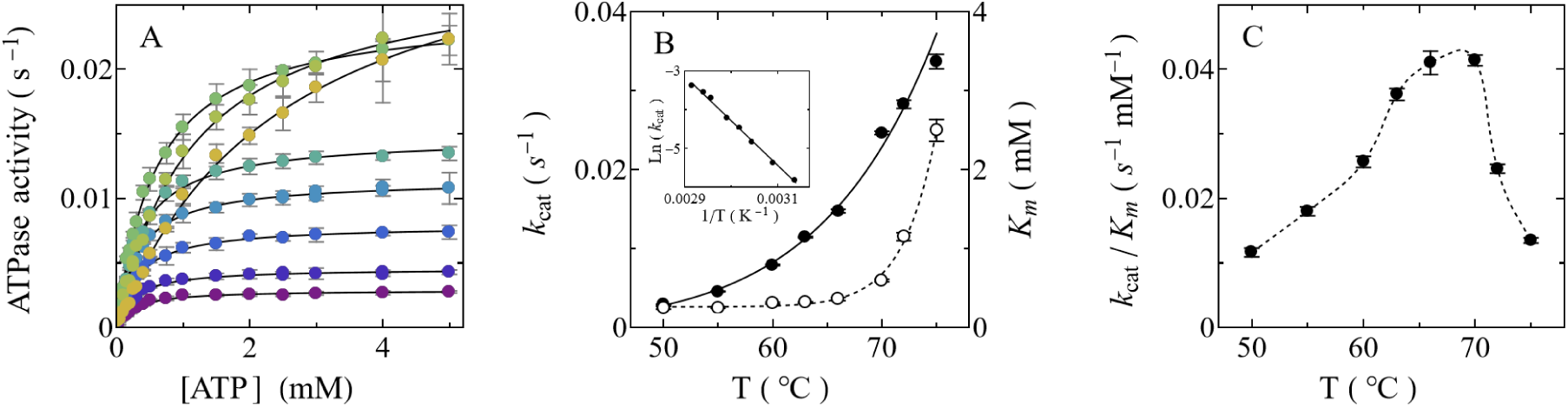
*Af*CopA-NP ATPase activity at different temperatures. ***A***. ATP concentration dependence of ATPase activity at different temperatures 75 (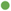), 72 (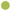), 70 (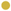), 66 (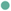), 63 (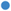), 60 (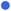), 55 (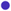), and 50°C (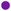), respectively. Continuous lines are the graphical representations of Equation 1 fitted to the experimental data of each curve. ***B.*** *Af*CopA-NP catalytic constant (*k*_cat_, 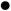) and Michaelis-Menten constant (*K_m_*, 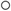) as a function of temperature. Continuous line is the graphical representation of the Arrhenius equation (Equation 2) fitted to the *k*_cat_ data. Dashed line constitutes a guide to the eye. Inset shows the corresponding Arrhenius plot. **C**. Catalytic efficiency (*k*_cat_/K_m_) as a function of temperature. Dashed line constitutes a guide to the eye.

It can be observed that ATPase activity follows a hyperbolic function of ATP concentration (Figure 1A):

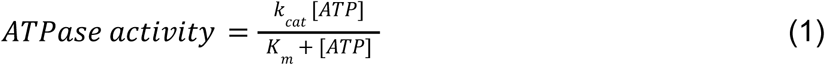

The catalytic constant (*k_cat_*) increases with temperature (Figure 1B) and this increase is well described by the Arrhenius equation.

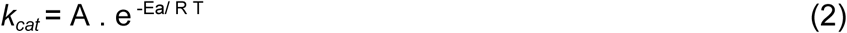

In the examined temperature range it does not evidence any curvature that would suggest a change in the catalytic mechanism. The activation energy (*E_a_*) obtained was 23.07 ± 0.02 kcal mol^-1^ and is approximately two-times the *E_a_*of the entire *Af*CopA ATPase (14 ± 1 kcal mol^-1^) ^46^. This finding is reflected in the *k_cat_* value observed for the isolated unit which is lower than the one corresponding to the full-length *Af*CopA ^48^. Additionally, the Michaelis-Menten constant (*K_m_)* also increases with temperature (Figure 1B), indicating a decrease of the apparent affinity for ATP. The different trends of the effect of temperature on *K_m_* and *k_cat_* yield a catalytic efficiency (*k_cat_/K_m_*) (Figure 1C) with a maximum around 70°C which is close to the observed optimal growth temperature of *Archaeoglobus fulgidus* ^49^. Notably, although the NP unit is in its isolated form, not only it remains catalytically active at the optimal working temperature of the full-length *Af*CopA as was previously mentioned ^21,22^, but also the apparent affinity for ATP (0.25 ± 0.04 mM) is conserved.

### Tryptophan fluorescence and far-UV molar ellipticity evidence a complex energy landscape for the native ensemble of conformations

To evaluate the functionally related conformational ensemble of *Af*CopA-NP we performed a temperature ramp experiment followed by tryptophan intrinsic fluorescence and far-UV ellipticity, both transitions being reversible. It is worth mentioning that *Af*CopA-NP has a single tryptophan residue located at position 576 (full-length *Af*CopA numbering) which is in the interface of the N and P domains of the catalytic unit (Figure 2A).

**Figure 2.**
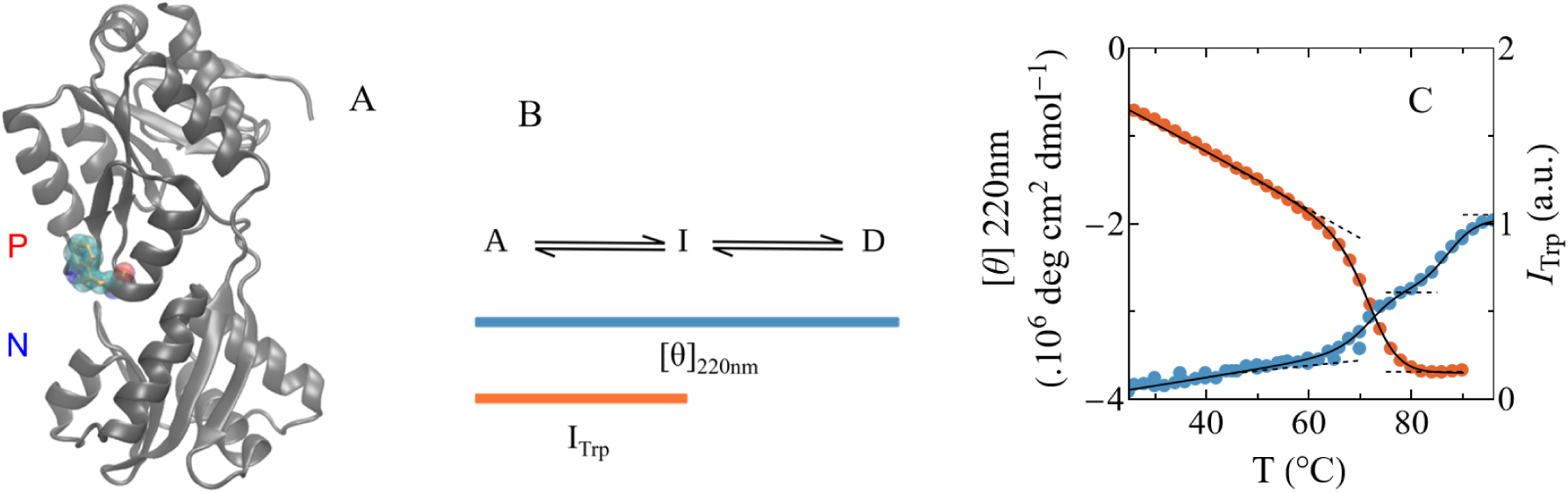
Spectroscopic analysis of *Af*CopA-NP at different temperatures. **A.** Structure representation of the PDB ID 3A1C chain A highlighting the tryptophan 576 of the *Af*CopA-NP (full-length *Af*CopA numbering) surrounded by its Van der Waals volume. Image was built using VMD software ^50^. **B.** Three-states equilibrium model. The only active state A is in equilibrium with non-functional species termed I (intermediate) and D (denatured). Color bars represent which transition is monitored with fluorescence (I_Trp_) or ellipticity ([θ]_220nm_). **C.** Intrinsic tryptophan fluorescence (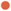) and far-UV molar ellipticity at 220 nm (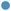) as a function of temperature. Continuous black lines are the graphical representations of the three-states model equation (Equation S2) fitted to both curves simultaneously (best fit parameter values are shown in Table S2). Dashed lines represent the pre- and post-transition slopes.

Altogether, temperature dependence of tryptophan fluorescence intensity and far-UV ellipticity were jointly described by the three states empirical model shown in Figure 2B (see AICc in Table S1 for comparing the fitting of Equations S1 and S2).The A to I transition would be the only one captured by fluorescence intensity because its Tm corresponds to the one with the lowest mid-denaturing temperature when monitoring ellipticity (T_m A-I_ = 72 °C, Table S2). The transition from I to D is only captured by far UV ellipticity and has a T_m I-D_ around 87 °C. Noteworthy, I is partially folded while D retains residual structure as revealed by the ellipticity content at 95 °C. The fact that temperature-denatured states retain residual structure has been previously documented in many proteins, being up to 50% of the original content ^51^.

### Small additions of urea enhances enzyme-catalyzed ATP hydrolysis suggesting a local unfolding mechanism related to catalysis

To monitor the link between ATP catalyzed hydrolysis and conformational rearrangements, we evaluated the effect of urea in *Af*CopA-NP activity at different temperatures (Figure 3). As expected, ATPase activity decreases with the addition of urea. However and surprisingly, at temperatures below the optimal working temperature ATPase activity first increases with the addition of low urea concentrations.

**Figure 3.**
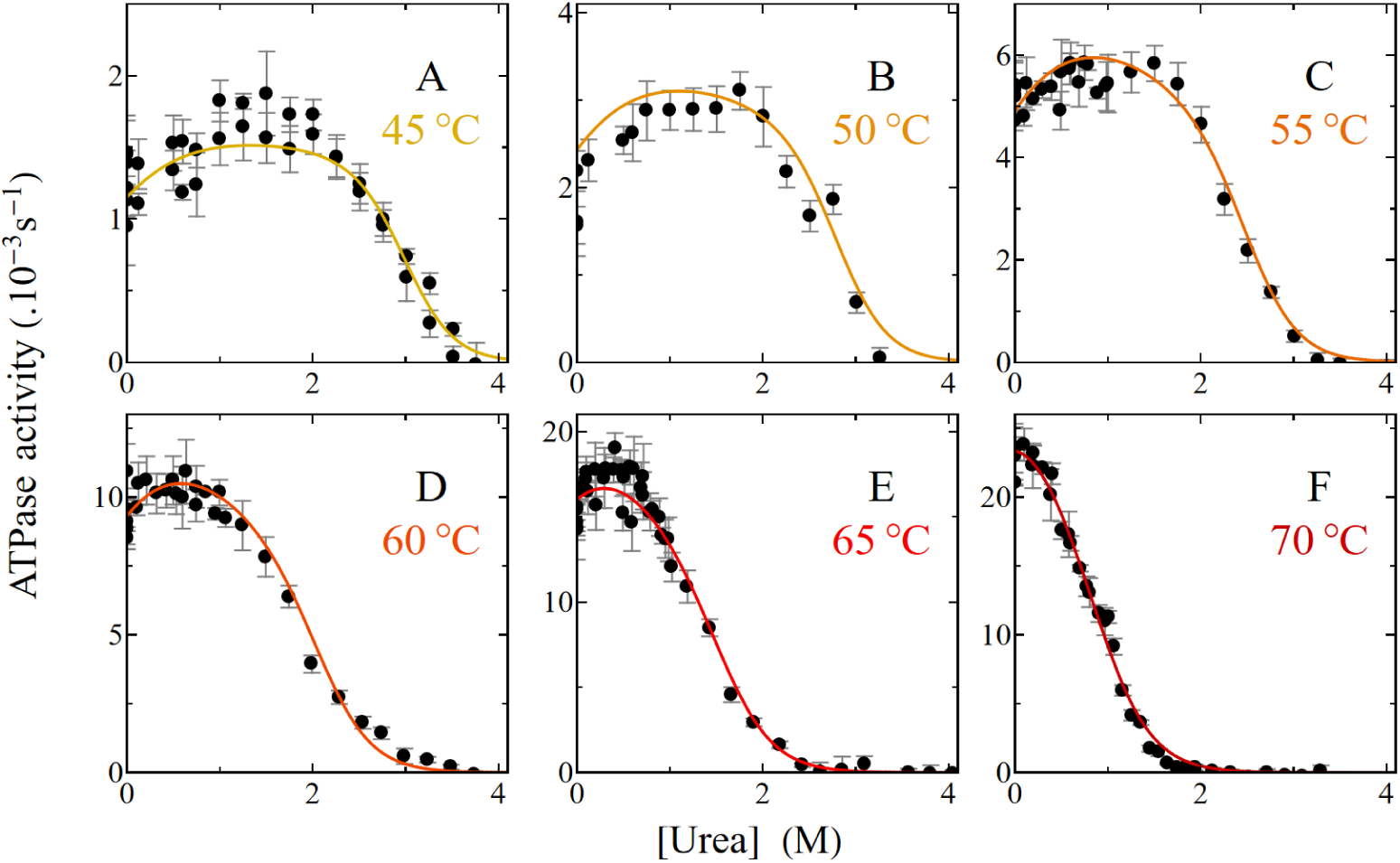
*Af*CopA-NP ATPase activity at different temperatures and urea concentrations. **A-F**. ATPase activity was determined at different temperatures (45, 50, 55, 60, 65, 70°C, respectively for each panel) in the presence of 3 mM ATP, 30 mM MgCl_2_ and 5 µM enzyme. The protein was incubated for at least 40 min prior to the reaction with the corresponding urea concentration. Continuous lines are the graphical representation of the equation derived from the model shown in Figure 9F (Equation S12) fitted to both spectroscopic and molar ATPase activity data at all temperatures, ATP and urea concentrations assayed. Best fit parameter values are shown in Table S3.

It can be observed that the urea concentration where the maximum ATPase activity is observed increases as temperature decreases.

Furthermore, in the presence or absence of urea, ATPase activity increases with ATP concentration following a hyperbolic function (Figure S3). The addition of urea leads to an increase in the apparent affinity (Km) and an increase in the *k*_cat_ at low urea concentrations (up to 2M). The temperature dependence of the *k*_cat_ in the absence and presence of low urea concentrations is shown in Figure 4.

**Figure 4.**
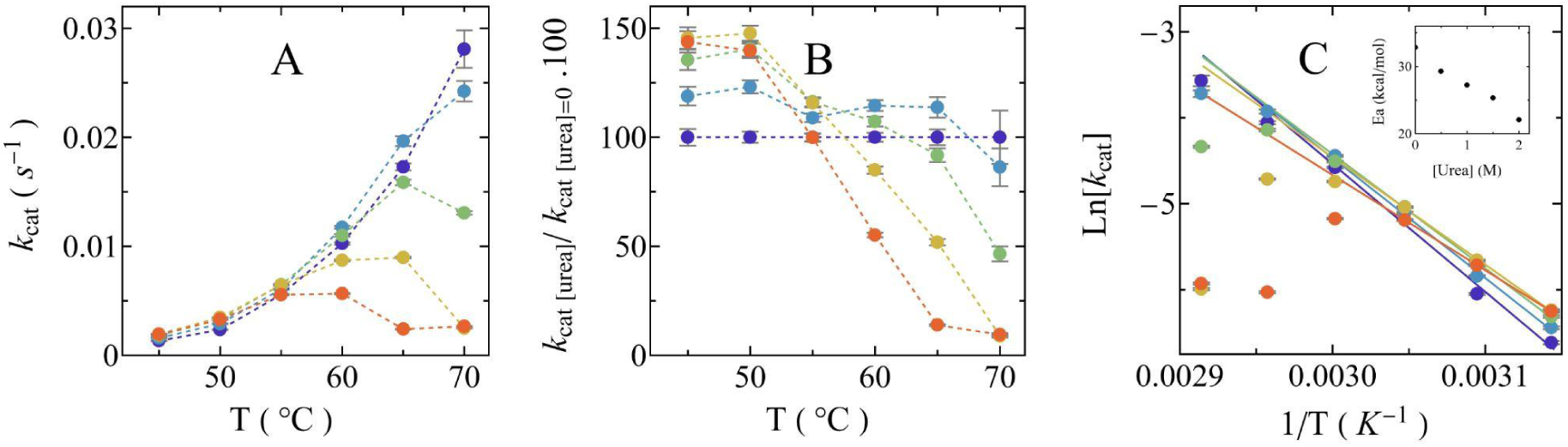
*Af*CopA-NP ATPase *k_cat_* at different temperatures and urea concentrations. **A.** *Af*CopA-NP ATPase *k_cat_* from Figure 3 shown at the representative values of 0 (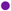), 0.5 (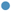), 1 (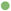), 1.5 (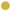) and 2 M urea (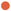) as a function of temperature. **B.** *k*_cat_ at the different urea concentrations relative to the value obtained in absence of urea ( *k*_cat [Urea]_ /*k*_cat [Urea]=0_ .100) as a function of temperature. **C.** Arrhenius plot of the *k_cat_* at the different urea concentrations. Lines are the graphical representations of the Arrhenius equation (Equation 2) fitted to the *k*_cat_ data as a function of the temperature. Fitting was performed considering only the data where, at all urea concentrations, *k*_cat_ increases with temperature (45 to 55°C). Inset shows the corresponding best fit values of E_a_ as a function of urea concentration. The coloring scheme is the same for the entire figure. In all cases, dashed lines constitute a guide for the eye.

The catalytic constant reaches its maximum at a specific temperature, and this temperature decreases upon urea addition (Figure 4A). When *k*_cat_ values are normalized to those in the absence of urea, it is evident that the catalytic constant increases with the addition of the chaotrope by up to 50% between 40 and 50°C (Figure 4B). Notably, while this enhancement occurs across all tested temperatures, the increase is most pronounced at the lowest temperatures assayed.

The increase of *k*_cat_ with temperature can be described, at each urea concentration, by the Arrhenius equation (Equation 2 and Figure 4C). It is noteworthy that the addition of small amounts of urea lowers the activation energy of the reaction (inset in Figure 4C). These results suggest that the rate limiting step of ATP catalysis is sensitive to small amounts of chaotropic agents. This can be analyzed in the context of the Corresponding States Hypothesis ^35^ which links protein activity and dynamics. When performing catalysis at the optimal working temperature protein dynamics are enough to achieve catalysis. However, as temperature decreases, these might be suboptimal. So the fact that the urea concentration corresponding to the maximum activity decreases with temperature seems feasible due to the effect of urea in protein flexibility and dynamics as addition of chaotropes favor local unfolding and may improve enzyme activity ^36,52^.

The conditions at which activation by urea occurs were also evaluated using tryptophan intrinsic fluorescence and far-UV ellipticity at different urea concentrations and temperatures (Figure 5).

**Figure 5.**
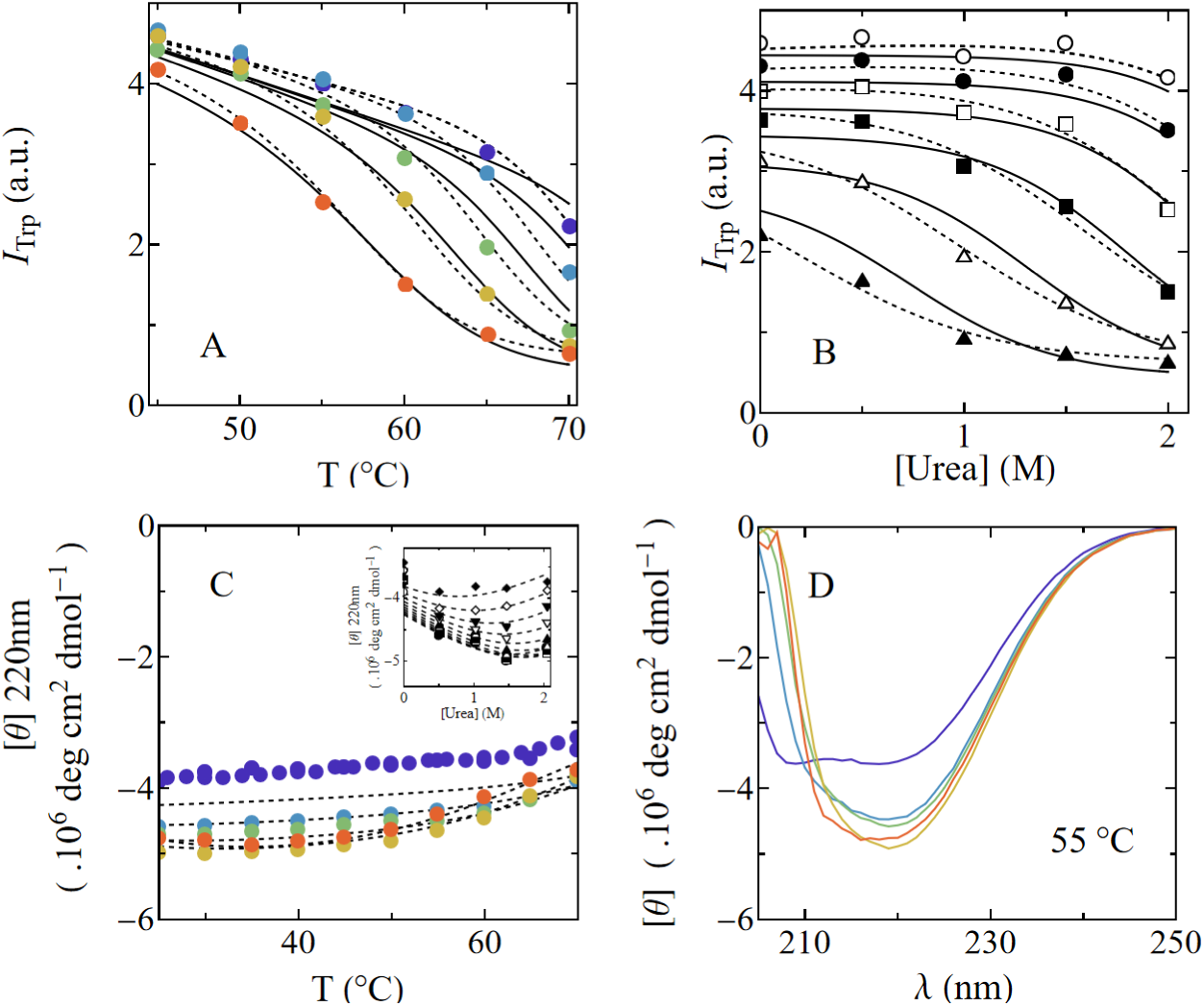
Spectroscopic analysis of *Af*CopA-NP at different temperatures and urea concentrations. **A.** Intrinsic tryptophan fluorescence as a function of temperature in the presence of 0 (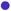), 0.5 (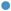), 1 (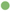), 1.5 (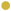) and 2 M urea (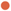). **B**. Intrinsic fluorescence intensity as a function of urea concentration at 45 (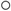), 50 (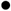), 55 (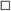), 60 (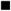), 65 (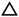) and 70 (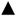) °C. In both panels A and B, dashed lines are the graphical representations of a two-states model (Equation S1) fitted to all curves simultaneously, with the best fit parameter values shown in Table S4. Continuous lines are the graphical representation of the equations derived from the model in Figure 9F (Equation S11) fitted to the experimental data. Best fit parameter values are shown in Table S3. **C.** Far-UV molar ellipticity at 220 nm as a function of temperature in the presence of the different urea concentrations. Inset shows the behavior as a function of urea concentration. Dashed lines constitute a guide for the eye. **D.** Far-UV ellipticity spectra at 55°C and the different urea concentrations. The coloring scheme is the same for the entire figure. When the measurements were performed in the presence of urea, the protein was incubated for at least 40 min prior to the reaction.

It is noteworthy that the conformational change evidenced by tryptophan fluorescence occurs at lower temperatures when small urea amounts are added (Figure 5A). Similarly, the same signal shifts to lower urea concentrations as temperature increases (Figure 5B). These changes are in line with the behavior of the maximum ATPase activity (Figures 3 and 4), suggesting a coupling between activity and this signal. Furthermore, when molar ellipticity at 220 nm was evaluated as a function of temperature (Figure 5C), there is no evidence of protein unfolding. Nevertheless, it is noteworthy that when urea concentrations as low as 0.5 M were added, there is a large reversible change in the far-UV ellipticity spectra as seen in Figure 5D.

The increase in catalytic activity in the presence of low denaturant concentrations has already been described for a small set of proteins including adenylate kinase, dihydrofolate reductase, prostaglandin D synthase, biliverdin-IXα reductase and papain ^36,52–61^. In general this phenomena is interpreted as the modulation of the transition barrier connecting functional conformations in a way that small additions of chaotropes lower its energy making the exchange between conformations more favorable. While theoretical studies of this mechanism are supported by experimental evidence, this evidence is largely limited to adenylate kinase ^52,55–57^ and, to a lesser extent, dihydrofolate reductase ^53,54^. Therefore, *Af*CopA-NP represents a new experimental model for exploring the connection between local unfolding and catalytic mechanisms.

### Tryptophan fluorescence captures conformational transitions that are linked to enzyme catalysis

As previously mentioned, the single tryptophan residue of *Af*CopA-NP is located at the interface between the N and P domains (Figure 2A) with a calculated surface exposed to the solvent of 50 % ^62^. This result is in agreement with the measured center of spectral mass of approximately 351 nm (at 25°C). When tryptophan fluorescence is evaluated as a function of urea concentration for a given temperature (i.e. 55°C, Figure 6A), total intensity abruptly changes, with a mid-denaturant concentration (C_m_) of 2.15 ± 0.03 M at 55°C.

**Figure 6.**
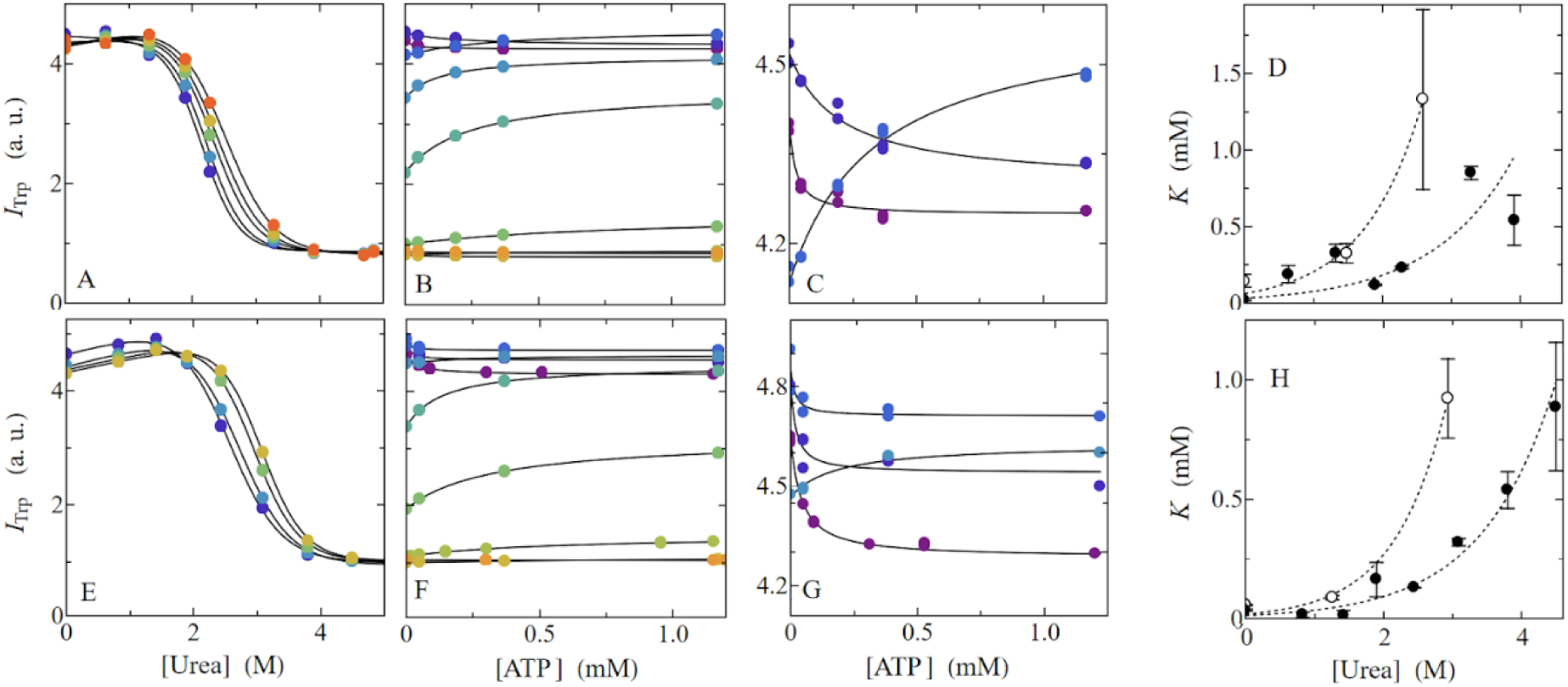
Effects of temperature and urea on the *Af*CopA-NP Intrinsic tryptophan fluorescence **A** and **E.** Intrinsic tryptophan fluorescence (I_Trp_) at different urea concentrations at 55°C (Panel A) and 0 (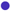), 0.05 (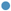), 0.2 (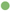), 0.4 (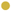) and 1.2 mM ATP (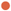) or at 45°C (Panel E) and 0 (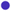), 0.05 (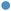), 0.3 (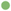) and 1.2 mM ATP (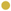). Continuous lines are the graphical representation of Equation S1 fitted to each dataset. **B** and **F**. Tryptophan fluorescence(I_Trp_) at different ATP concentrations at 55°C (Panel B) and 0 (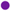), 0.6 (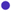), 1.3 (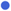), 1.9 (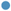), 2.3 (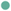), 3.3 (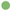), 4.0 (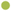) and 4.8 M urea (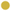) or at 45°C and 0 (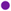), 0.83 (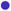), 1.42 (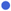), 1.9 (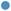), 2.44 (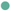), 3.09 (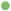), 3.8 (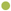), 4.5 (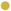) and 5.3 M urea (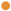). Continuous lines are the graphical representation of Equation S5 fitted to each dataset. **C** and **G**. A zoom of panels B and F shows the corresponding behavior at the lowest urea concentrations. **D** and **H**. Dissociation constant *K*_ATP_ values (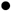) obtained from the best fit of Equation S5 to _Trp_ dependence on ATP concentration at 55 and 45°C respectively. Panels also show ATPase activity *K*_m_ values (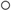) obtained from the best fit of Equation 1 to ATPase activity dependence on ATP concentration as a function of urea concentration at the same temperatures (Figure S3). Dashed lines are the graphical representation of exponential functions fitted to each dataset.

The center of spectral mass also changes abruptly, from approximately 351 nm to 356 nm (Figure S4), indicating an increase in solvent exposure. Addition of ATP shifts C_m_ values towards higher urea concentrations (Figure 6A and S4), evidencing ATP binding to *Af*CopA-NP from this protecting effect. Interestingly, ATP exerts a dual effect on fluorescence intensity. While at low concentrations it quenches the emission, at higher concentrations it enhances it (Figures 6B and 6C), suggesting the existence of an equilibrium of species with different quantum yields. This overall behaviour is maintained at both the evaluated temperatures (Figures 6E-G). The results suggest that fluorescence is monitoring at least two partially independent phenomena: the relative movement between N and P domains and ATP binding, which takes place in the N domain as has been previously characterized for the human CopA homologue ^63^. Considering binding of a single ATP molecule per protein, at each urea concentration, apparent affinity for ATP (*K*_ATP_) can be obtained using Equation S5. As can be seen in Figures 6D and 6H, *K*_ATP_ increases with urea concentration.

### Local frustration is linked to open-closed transition

As discussed by *Jayakanthan et al*., ATP binding to the catalytic unit of most P-type ATPases induces a rotation of both N and P domains leading to a more compact conformation and resulting in a decrease of the binding site solvent exposure ^23^. Structural studies of *Af*CopA-NP have shown only closed conformations (Figure S5) ^21,22^. However, it is reasonable to presume the existence of an open state which has been reported to be relevant to the catalytic mechanism ^23^. Energy landscape theory provides a thorough framework for analyzing protein folding and conformational transitions ^64^. These transitions occur in a funneled landscape where roughness appears as patches of highly frustrated contacts which are usually involved in conformational transitions and functional dynamics ^65^. Frustration patches in the closed structure of *Af*CopA-NP, in the presence of an ATP analogue (PDB ID 3A1C), are mainly located at the interface between N and P domains (Figures 7A and 7C). To assess whether these patches are involved in conformational transitions, we followed the protocol implemented by Guan et al ^42^. Briefly, after identifying highly frustrated positions in the closed structure we replaced those positions in the multiple sequence alignment provided to AlphaFold2 with gaps (provided as a Supplementary list - List S1). The top scored prediction revealed a conformation where the *Af*CopA-NP binding site is exposed to the solvent (Figures 7B).

**Figure 7.**
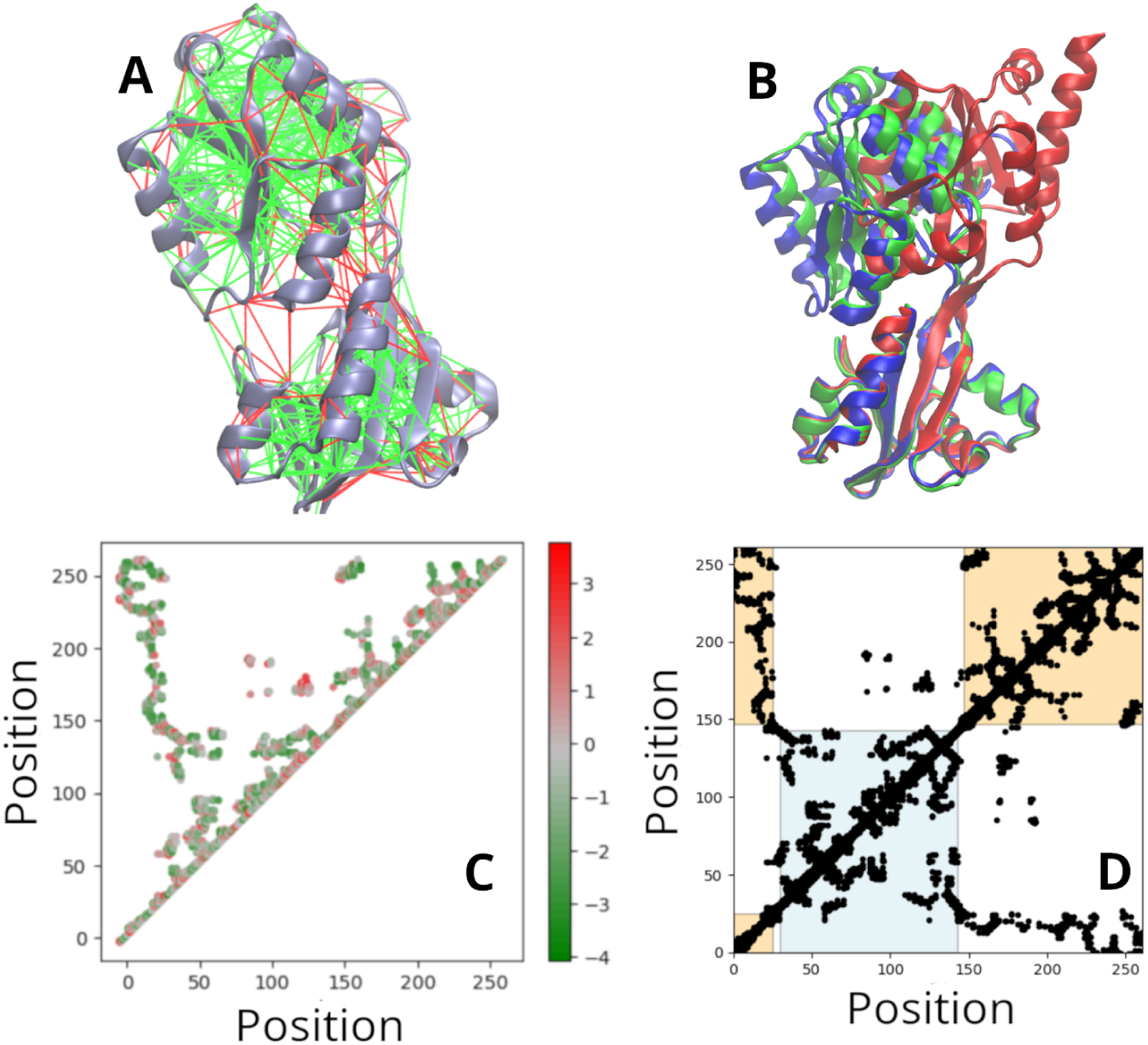
Effect of frustration in the shape of the conformational ensemble of *Af*CopA-NP. **A**. Mutationally frustrated contacts were calculated as described in Materials and methods using the Frustratometer server^66^ and are displayed on *Af*CopA-NP PDB ID 3A1C structure. Red and green lines correspond to the highly and minimally frustrated contacts, respectively. **B**. Experimentally reported structure with ligand-analog bound (PDB ID 3A1C) (blue), in the absence of the ligand (PDB ID 2B8E) (green) and A2F predicted structure in the absence of ligand (red). **C.** Contact map colored by mutational frustration index. Each point corresponds to residues that are in contact while the color scale corresponds to the local frustration index. Position numbering is according to the *Af*CopA-NP sequence. **D.** Contact map colored per domain contacts. In light blue are the contacts of the N domain, in orange contacts of the P domain, and uncolored are the inter domain contacts.

To characterize the transition between both conformations, we performed structure-based α-carbon simulations using an in-house implementation of the forcefield described in *Clementi et al*. ^67^. To explore the dynamics of the open-closed rearrangement we performed umbrella sampling guided simulations using the difference in the RMSD between the closed (PDB ID 3A1C) and the AF2/Local frustration-predicted open structure (Figure 7B, red) as the reaction coordinate (RMSD_difference,_, Equation 4). We analyzed the results considering the contacts that are formed exclusively in the closed conformation (Q_ligand_) and RMSD_differnce_ coordinates. Varying the energy weight of the Q_ligand_ (ε_Qligand_) populates the open (ε_Qligand_ = 0) or the closed state (ε_Qligand_ = 1). Free energy profiles with different ε_Qligand_ values are shown in Figure S6. When ε_Qligand_ = 0.8 the resulting free energy landscape shows three main basins being the open (low Q_ligand_ and low RMSD_d_), a partially closed (intermediate Q_ligand_ and high RMSD_d_) and a closed (high Q_ligand_ and high RMSD_d_) states (Figure 8A). Interestingly, the partially closed conformation appears with a slight disruption of ligand-mediated contacts (Figure 8B and Equation S13).

**Figure 8.**
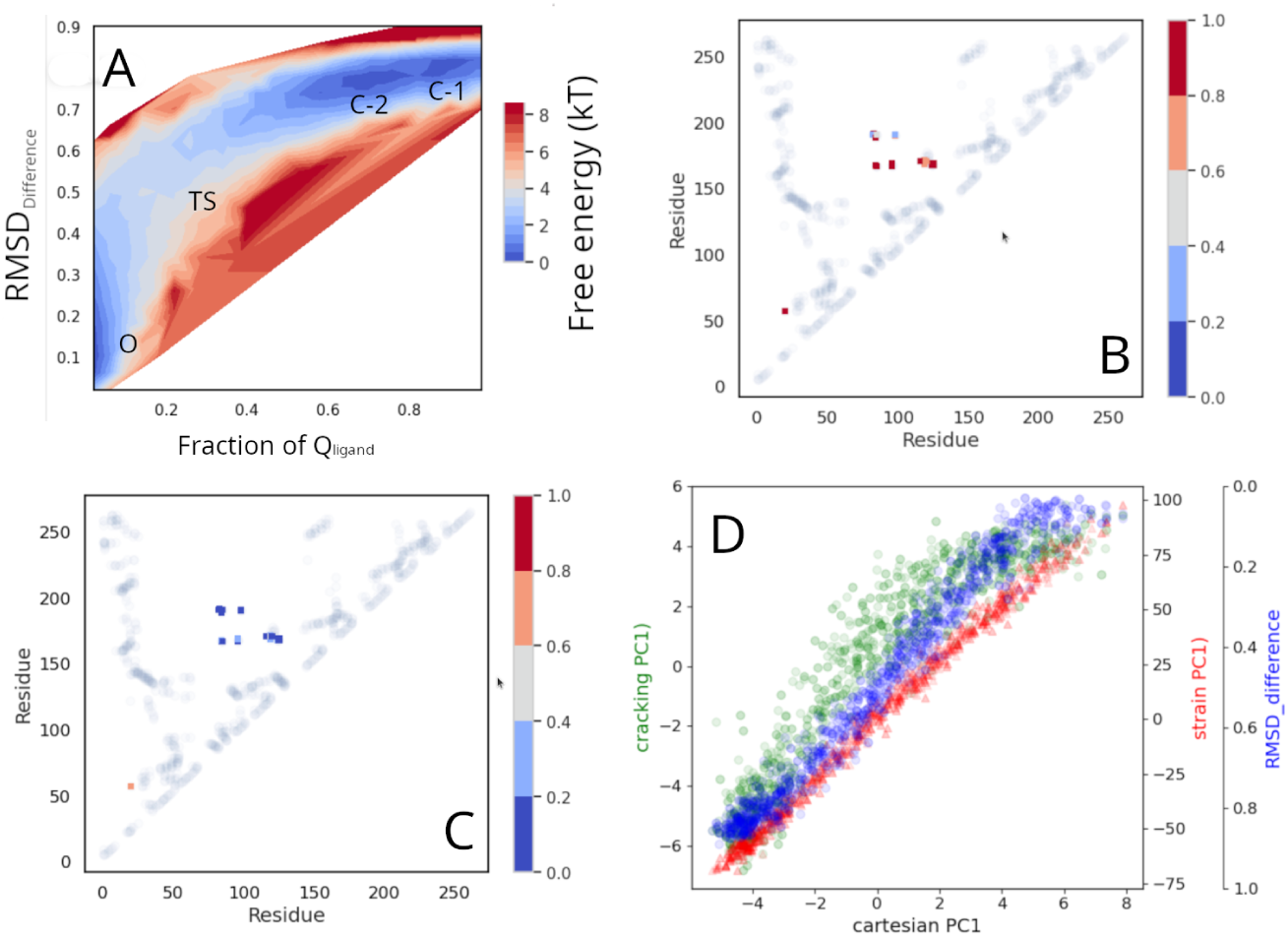
Energy Landscape of the open-closed transition of *Af*CopA-NP. **A**. Free Energy profile using the fraction of Q_ligand_ and RMSD_difference_ as reaction coordinates at ε_Qligand_ = 0.8. The color represents the free energy in kT units. Labels indicate open (O), transition (TS), partially closed (C-2) and closed conformations (C-1). **B**. Contact α-values for the C-2 conformation (Equation S13). **C**. Contact α-values for the TS (Equation S13). Position numbering in panels B and C is according to the *Af*CopA-NP sequence. **D**. Correlation of the projections of the PC1 of cracking (green), first Principal Component of strain (red) and RMSD_difference_ (blue) with the projection of the cartesian first principal component to the trajectory.

This result implies that destabilizing those contacts favors a partially closed state prior to the opening transition. This may be mimicking the addition of urea which at below denaturing concentrations can destabilize the already formed contacts. The transition state, on the contrary, has most of the Q_ligand_ disrupted as can be seen in Figure 8C. The addition of urea at small concentrations then would favour the partially closed conformation making the opening transition more favorable. A picture of the main ensemble of conformations can be seen in Figure S7.

To analyze the conformational change we calculated the strain and cracking principal components as described in Materials and methods ^42^. The largest-amplitude principal component (PC1) on interatomic distances (strain) and formation or rupture of native contacts (cracking) show that the involved contacts are those with highest frustration index (Figures S8 and 7C). Figure 8D shows a perfect correlation between the PC1 both the cartesian coordinates, strain and RMSD_difference_ (blue dots). This indicates that these components are describing the same transition. The increase in the strain PC1 also correlates with the opening of *Af*CopA-NP (low values of RMSD_difference_) indicating a stressing Q_ligand_ interface. It can also be seen that the cracking PC1 (green dots) stops increasing after a certain value which reveals the breakage of Q_ligands_. All these results point to a cracking mechanism linked to the functional movements. The modulation of ε_Qligand_ would be mimicking the urea effect.

### ATPase activation is a consequence of the accumulation of a reaction intermediate

To gain insights into the coupling between ATP hydrolysis and conformational changes we analyzed both the spectroscopic and activity results in terms of a model that comprises two states, I and A, where only A is able to bind ATP and catalyze its hydrolysis (Figure 9).

**Figure 9.**
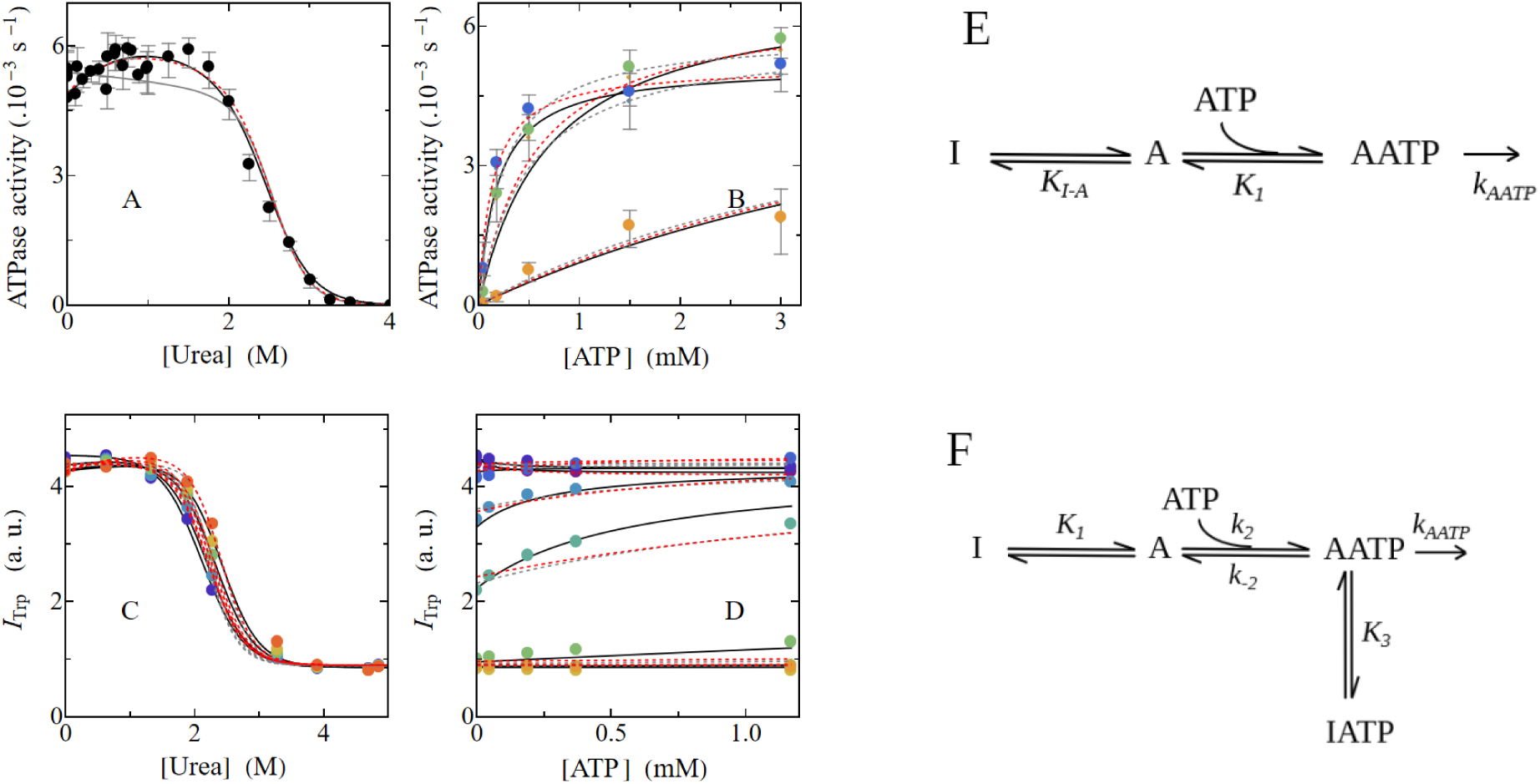
Description of the effects of ATP and urea on ATPase activity and intrinsic tryptophan fluorescence at 55°C. **A-D**. Gray and red lines are the graphical representations of equations derived from model in panels E (Equations S6 and S7) and F (Equations S8 and S9) globally fitted to the experimental data previously shown in Figures 3C, 6A-D and S3B, both under the rapid equilibrium assumption. Black lines are the graphical representations of the equations derived from Panel F without assuming rapid equilibrium conditions (Equations S8 and S10), globally fitted to the experimental data. **E**. Reaction model of *Af*CopA-NP catalytic activity with only one ATP binding step. A is in equilibrium with a reaction intermediate state termed I and is able to bind ATP to give AATP, which is the catalytically active species. **F**. Reaction model of *Af*CopA-NP catalytic activity with two ATP bound species but only one catalytically active. The state A is in equilibrium with a reaction intermediate termed I. When the model is solved under the assumption of rapid equilibrium *k*_-2_/*k*_2_ is replaced by the equilibrium constant *K*_2_. State A is able to bind ATP and populate two species, AATP and IATP, where only AATP is active. Table S5 shows the AICc values calculated with Equation 3, corresponding to the fitting of the three models previously described and the best fit parameter values corresponding to the best model are shown in Table S6.

Although this model (represented by Equations S6 and S7) is able to explain the change of intrinsic fluorescence with urea and ATP, the activation exerted by the addition of small urea amounts is not captured (Figure 9, gray lines). Moreover, if we follow the approach described by Rogne et al. ^52^ making *k*_AATP_ dependent on urea concentration, we can explain the ATPase activity increase but we fail to describe the intrinsic fluorescence change and the apparent affinity for ATP (Figure 9, red lines). Alternatively, we developed a model (Figure 9F) that incorporates one additional ATP bound species, IATP, in equilibrium with AATP, with only the latter catalytically active. The model was solved under rapid-equilibrium assumption for all steps (K_2_=*k*_-2_/*k*_2_, Equations S8 and S9). Interestingly, although this model is able to explain the dependences of both ATPase activity and Trp fluorescence on urea concentration, it does not fully explain the dependences of both signals on ATP concentration. However, when the ATP binding step is considered out of the rapid equilibrium condition (Equation S10) all the data are fully described (Figure 9 solid lines for 55°C and Figure S9 for 45°C). Notably, the fact that this step is out of the rapid equilibrium condition yields a difference in *K*_m_ and *K*_ATP_ dependence on urea concentration (Figure S10).

Figure S11 shows the molar fractions of each species at different temperatures, urea and ATP concentrations. The activating effect at low urea concentrations would be exerted by an increase of the concentration of species AATP with respect to IATP. The equilibrium among these species and their slightly different quantum yield, account for the effect observed on tryptophan fluorescence. At higher urea concentrations, dissociation of ATP is favoured, with the uppermost displacement of the equilibrium between A and I, giving place to the decrease in ATPase activity and the total intensity fluorescence. Interestingly, the *m-values* (Table S6), which are the dependence of each constant with the urea concentration, are compatible with partial unfolding reactions (A to I in Figure 9F). A small *m-value* corresponding to ATP binding (*m*_2_, A to AATP) suggests a small change of accessible surface upon binding. In contrast, the *m-value* when moving from IATP to AATP (*m*_3_) suggests that the productive AATP state is less compact than IATP. We have shown in Figure 5D that upon addition of small amounts of urea there is a large and reversible change in secondary structure which is compatible with stabilization of beta sheets and destabilization of helical motifs. This change takes place in the same range of urea concentrations where activation occurs.

Furthermore, we evaluated whether this model was also able to describe the temperature dependent observations. Temperature dependence of the step governed by *K_1_* required inclusion of the heat capacity change to fully describe our results. This is reasonable given the large change in accessible surface area associated to that step (*m*_1_ in Table S6). When considering the effect of temperature, the model not only describes ATPase activity dependence on urea concentration (continuous lines in Figure 3, Figure S12) at all assayed temperatures, but also predicts tryptophan fluorescence dependence on both temperature and urea concentration (continuous lines in Figures 5A and 5B, Figure S12). The fact that the urea activating effect is lower at the highest temperatures can now be explained by decrease in the accumulation of AATP as the temperature increases (Figure S13). The best fit value of *E_a_* of the catalytic step (31 kcal mol^-1^) is of the order of that obtained from the dependence of *k*_cat_ with temperature (Figure 1B - 23kcal mol^-1^) which reinforces the robustness of our modelistic description of the reaction.

### The conformational changes of the isolated catalytic unit are part of the P-type ATPases reaction cycle

As described in the introduction, P-type ATPases reaction cycle is characterized by the presence of a phosphorylated intermediate and can be described by the Post-Albers model. Briefly, this model includes two main conformational ensembles, E1 and E2, ATP binding, hydrolysis, autophosphorylation and ion transport (Figure S1) ^3,10^. In this work we studied the ATP hydrolysis reaction performed by the catalytic unit, thus, the mechanism described is a part of the Post-Albers mechanism.

In the case of full length Cu(I) ATPases, even though structural information is still scarce to characterize the Post-Albers cycle completely, we evaluated the conformations of the NP unit in it (Figure 10, structures in gray). Structures in E1 conformation were reported for the *Archaeoglobus fulgidus* variant either in the absence or in the presence of Cu^+^ (PDB IDs 7R0G and 7R0H, respectively) ^47^ and the *Legionella pneumophila* variant *Lp*CopA in the absence of ligands (PDB ID 3RFU) ^68^. In these structures, the catalytic unit shows different conformations. In *Lp*CopA (PDB ID 3RFU), it is in an open conformation, similar to the model we obtained using AlphaFold2 (Figure S5B). On the contrary, in *Af*CopA in the absence of ligands it is in a closed conformation (PDB ID 7R0G) (Figure 10 and S5A). Moreover, in *Af*CopA in the presence of Cu(I) (PDB ID 7R0H) it is in a partially closed conformation that is similar to the reported structure of the isolated *Af*CopA-NP from PDB ID 2B8E (RMSD 0.85 Å) ^21^. Interestingly, the catalytic unit reported in PDB 2B8E is consistent with the partially closed ensemble, denoted as C-2 in Figures 8 and S7, obtained from the structure-based model simulations performed in this work. These conformations seem to facilitate ATP binding. Following the reaction cycle, no structures for the full length protein in E1ATP conformation were reported, but the structure of the isolated *Af*CopA-NP in the presence of an ATP analogue (PDB ID 3A1C) ^22^ shows a closed conformation. After ATP binding, the enzyme is phosphorylated forming E1P-ADP, with no experimental structures associated. ADP is then released, the phosphorylated E1P shifts to E2P and transitions to E2-Pi (PDB IDs 4BBJ and 4BYG for E2 and E2Pi, respectively) ^20,69^ with both P and N domains interacting with A domain. In these structures the catalytic unit is in an open conformation, comparable to our predicted *Af*CopA-NP open structure. Taking all this information into account, we hypothesize that the enzyme in E1 is in equilibrium between conformations where the catalytic unit is in open and closed states. Binding of Cu(I) induces a partially closed conformation, which could facilitate ATP binding. After phosphorylation, A domain interacts with the catalytic unit promoting its opening, as shown in the structures of E2P and E2-P. Finally, Pi release would leave the enzyme capable of initiating a new turnover cycle.

**Figure 10.**
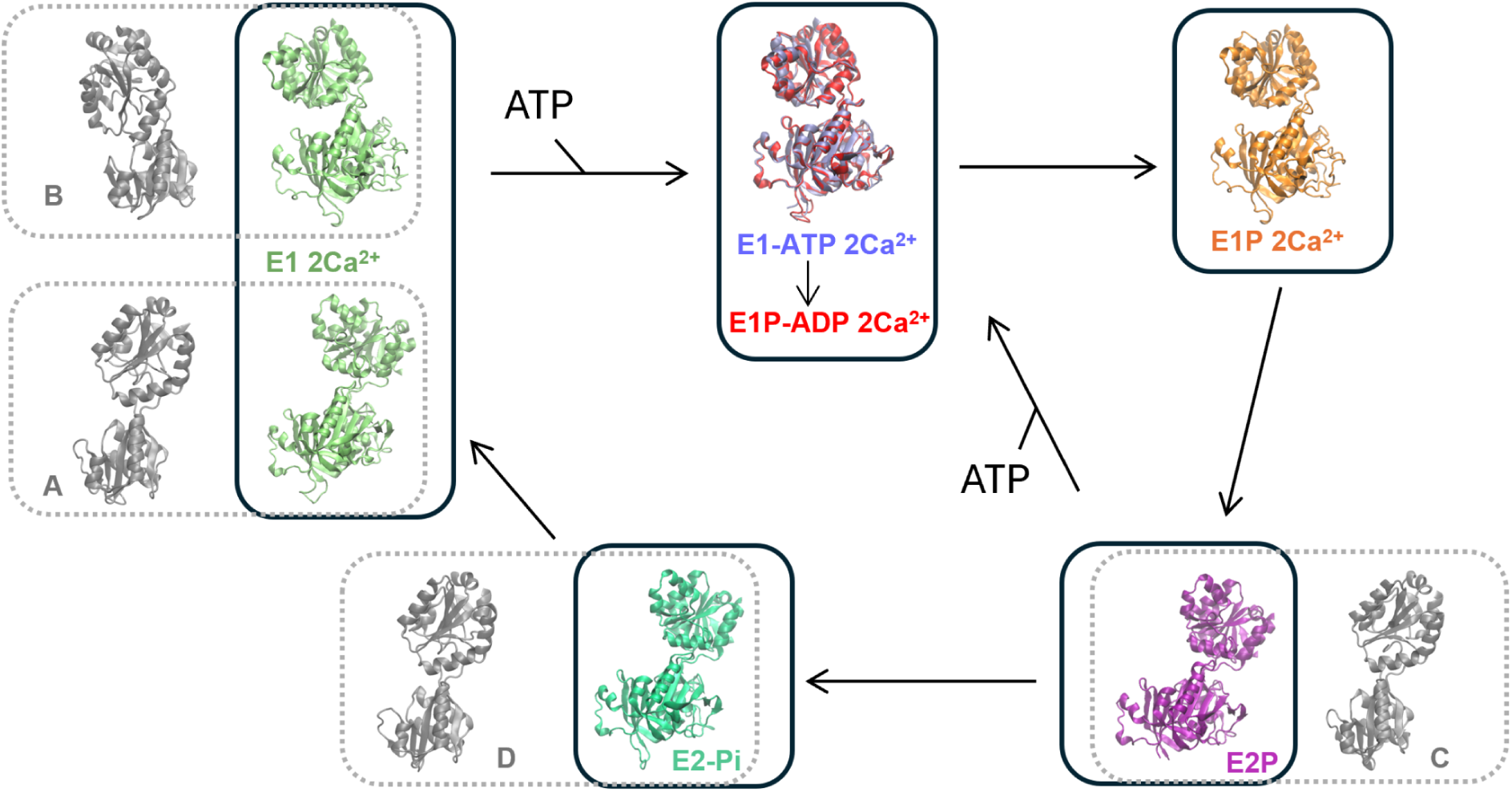
Catalytic unit conformations along the Albers-Post model. Inside each box is shown the structure of the catalytic unit of SERCA at the corresponding reaction steps. E1 2Ca^2+^ conformation was described in two conformations (PDB ID 1SU4 on the bottom and PDB 7E7S on the top). The structures for the rest of the conformations are shown in the corresponding color and consist of E1-ATP 2Ca^2+^, E1P-ADP 2Ca^2^, E1P 2Ca^2^, E2P and E2-Pi (PDB IDs 6LLE, 1T5T, 7W7T, 6LLY and 7W7W, respectively). In gray are represented available structures for the catalytic unit of Cu(I)-ATPases: *Lp*CopA and *Af*CopA in the absence of ligands (A and B, PDB IDs 3RFU and 7R0H, respectively) and *Lp*CopA in a the presence of ligands (C and D, PDB IDs 4BBJ and 4BYG, respectively).

Although from a different subfamily, the structural information of the P_2B_-type Sarcoplasmic Reticulum Ca^2+^-ATPase allows a more detailed interpretation of the reaction cycle (Figure 10). In this case, the catalytic unit conformation is defined by the orientation of the helices 7 and 10 (similar to helices 5 and 7 from Cu(I) ATPases). Binding of calcium to E1 (E1 2Ca^2+^) leads to conformations with two different reported structures proposed to be in equilibrium: an open (PDB ID 1SU4) and a partially closed one (PDB ID 7E7S, similar to PDB ID 7R0H of *Af*CopA) ^30,70^. When the ATP analogue AMPPCP is bound, the catalytic unit is closed (E1-ATP 2Ca^2+^, PDB ID 6LLE) supporting our observations on the isolated unit of *Af*CopA-NP in the presence of the same analogue ^70^. The closed conformation is maintained even after enzyme phosphorylation (E1P-ADP 2Ca^2+^, PDB ID 1T5T) ^71^. When ADP is released the catalytic unit turns to a partially closed conformation (E1P 2Ca^2+^ PDB ID 7W7T) and then opens (E2P, PDB ID 6LLY) ^72,73^. In this conformation, the A domain interacts both with N and P domains. Dephosphorylation leads to E2-Pi conformation (PDB ID 7W7W), where the catalytic unit is open as discussed for the *Lp*CopA ^72^. Furthermore, as it was described for SERCA, ATP can bind to the enzyme in both of the E1 2Ca^2+^ equilibrium conformations: open or partially closed ^70^. Moreover, ATP can also bind to E2P, enhancing enzyme activity with a modulatory effect on the reaction cycle ^74^.

In our work, we explored the link between conformational rearrangements of the catalytic unit and its ability to perform catalysis. We showed that catalysis is modulated by ATP-bound active and inactive species, and that urea induces an increase in the catalytic efficiency by influencing the equilibrium between these species. However, we are aware that urea does not play a physiological role in the catalytic cycle of these P-type ATPases. Thus, how could cracking be relevant for the protein function?. In the context of the available structures of the Post-Albers cycle, E1-ATP represents an ensemble of conformations where A and N domains are interacting, forming a binary complex. Also, E2P conformations involve the interaction of A domain with the phosphorylated aspartic acid in the P domain from the catalytic unit forming a ternary complex. Furthermore, the transition from E1-ATP to E2P ensemble occurs with breakage of contacts involving the interface between N and P domains. On each catalytic cycle, the N domain disengages from the P domain, allowing A domain to associate with it. This exchange is tightly synchronized with subsequent triggering events—such as substrate binding or product release—to ensure continuity in ATP hydrolysis. This contact disruption mediated by the A domain, resembles what urea is doing in our experiments by disruption of the mentioned interface. A comparable mechanism has been observed in the genetical regulation network lκβ/Nκβ/DNA, where the transcription factor Nκβ initially binds DNA to form a binary complex and is subsequently displaced by lκβ through the formation of a transient ternary intermediate ^75^. This phenomenon, known as *molecular stripping*, has also been documented in various protein-protein interactions ^76–78^. Such mechanism enables kinetic regulation of binding and dissociation events, challenging the traditional view that relies solely on thermodynamic control. Although hypothetical, the disruption of the NP interface upon A domain rotation may be occurring through this type of mechanism, having a consequence in enzymatic catalysis.

## Concluding remarks

Our results highlight the intricate regulation of *Af*CopA-NP ATPase activity by temperature and small amounts of urea. The temperature-dependent changes in ATP apparent affinity (*K*_m_) and catalytic turnover (*k_ca_*_t_) resulted in a biphasic catalytic efficiency profile, with an optimal temperature of approximately 70°C. The linearity of the Arrhenius plot across all conditions points to a consistent reaction mechanism, despite variations in temperature and urea concentration. The addition of small amounts of urea enhanced ATPase activity, contrasting with its typical denaturing effects. Spectroscopic analyses evidenced a strong link between conformational transitions and enzyme catalysis showing that addition of ATP significantly affects intrinsic tryptophan fluorescence, and in the presence of urea, reveals the presence of distinct ATP-bound species in equilibrium. Interestingly, complementary far-UV ellipticity measurements revealed that the transition observed by fluorescence spans only a partial unfolding event as revealed by the high ellipticity content. Altogether, these results evidence an intricate relationship between conformational dynamics and enzyme catalysis. This provides valuable insights of the functional energy landscape and a role of local unfolding that facilitates catalysis.

Local frustration analysis revealed that the interface between the nucleotide binding and phosphorylation domains is enriched in highly frustrated contacts. Combination of this information with AlphaFold2 allowed us to predict the *A*fCopA-NP open conformation and study the open-closed transition using structure-based model simulations, supporting the crucial role of local frustration in conformational rearrangements. As a result of these simulations, we found a partially closed conformation similar to the experimental structure of *Af*CopA-NP in the absence of ligands (PDB ID 2B8E). The fact that the highly frustrated interface in the closed conformation is partially disrupted suggests that this partially closed conformation may be populated when urea is added having a consequence in enzymatic catalysis. Thus, the observed effect on the enzyme activity may be the consequence of a cracking-like mechanism underscoring a delicate balance of structural and energetic factors driving enzyme function and regulation.

In summary, this work provides a comprehensive understanding of how temperature, urea, and conformational dynamics modulate *Af*CopA-NP catalytic activity. By integrating our experimental observations into a unified mechanistic model, we offer an explanation for the observed phenomena highlighting the critical role of conformational dynamics, local frustration, and “cracking” in enzyme regulation, obtaining crucial insights into the mechanistic principles governing protein function.

## Materials and methods

### Materials

Yeast extract and soybean tryptone were purchased from Britania (Buenos Aires, Argentina). Ampicillin was purchased from BioBasic (Toronto, Canada). Lactose, Tris and Imidazole were purchased from Anedra (Buenos Aires, Argentina). Buffer MES, NaCl, ATP and MgCl_2_ were from Merck (Darmstadt, Germany). Urea was purchased from MPbiomedichals (California, USA).

### Cloning of *Af*CopA-NP

*Af*CopA-NP (from K407 to K671, full length *Af*CopA numbering ^46^) was cloned into the pPR-IBA1 vector (IBA). The resulting construction (274 residues) contained a small linker followed by a 6xHis tag at the C-terminus of the protein ^21^. Cloning was confirmed by DNA sequencing of the plasmid amplicon.

### Strains, media and growth conditions

*Escherichia coli Bl21 Star (DE3) pLysS* strain was used for protein expression. Bacteria were grown at 37 °C in Luria Bertani broth (LB: 1% NaCl, 1% tryptone, and 0.5% yeast extract), supplemented with 100 μg/mL Ampicillin. Briefly, 25 ml LB media were inoculated with stock *Bl21 Star (DE3) pLysS Af*CopA-NP bacteria overnight at 37 °C with constant shaking. The culture was then diluted up to 0.01 absorbance units in 1L fresh LB media and growth at 37 °C with constant shaking up 0.6-0.8 absorbance units. At this point protein expression was induced by the addition of 10% Lactose and incubation continued for 3.5 hours. Cells were harvested by centrifugation and frozen at −80 °C until protein purification.

### *Af*CopA-NP purification

Cells were resuspended in 50 mM Tris pH 7.5 at 4°C and 500 mM NaCl (buffer A) plus 10 mM Imidazole and 1 mM phenylmethylsulfonyl fluoride. Lysis was performed by sonication using 8 pulses of 30 seconds separated by 1 minute of sample rest at 4 °C. Afterwards the suspension was sedimented for 15 minutes at 8.000*g* and 4°C. Supernatant was heated for 15 minutes at 55°C and centrifuged again for 30 minutes at 10000*g* for precipitation of thermolabile proteins as described in ^22^. The resulting supernatant was loaded in a Ni-NTA column and washed with 50 column volumes of Buffer A plus 20 mM Imidazole. Elution was done with buffer A plus 300 mM imidazole and followed by intrinsic tryptophan fluorescence. Fractions containing the protein were pooled and Imidazole was removed by overnight dialysis or passing the sample through a chromatographic column containing Sephadex G-25 equilibrated with storage solution (50 mM Tris pH 7,5 and 150 mM NaCl) and stored at −80 °C. Protein purity was evaluated by SDS-PAGE and its concentration was estimated by light absorbance at 280 nm (ε_280nm_ 5500 M^-1^cm^-1^) ^79^.

### Fluorescence measurements

Steady-state intrinsic tryptophan fluorescence measurements were performed in a Jasco FP-6500 spectrofluorometer equipped with a Peltier thermostatic cell holder. To this end, a 0.3 cm path length cell sealed with a Teflon cap was used. The excitation wavelength was 295 nm and emission data was collected in the range of 305–430 nm. The spectral slit-width was set to 3 nm for both monochromators. I_Trp_ was calculated as the total emission intensity. Results shown are representative of at least 3 independent experiments.

### Circular dichroism spectroscopy

Ellipticity of protein samples was evaluated using a Applied Photophysics Chirascan V100 spectropolarimeter equipped with a Peltier thermostatic cell holder. Far-UV circular dichroism spectra were recorded in the range between 205 and 250 nm, protein concentration was 10 μM, and a cell of 1 mm path-length was used. Data was acquired at a scan speed of 4 nm s^-1^. Blank scans were subtracted from the spectra and molar ellipticity was expressed in units of deg cm^2^ dmol^−1^.

### Protein unfolding experiments

After incubation for at least 40 min at room temperature, *Af*CopA-NP isothermal unfolding experiments were carried out in 0–2 M urea in a buffer solution (25 mM MES pH 5.5 at 70°C, 100 mM NaCl). The process was followed by far-UV ellipticity and intrinsic tryptophan fluorescence. Thermal unfolding experiments were performed by varying the temperature from 25 to 70°C or 96°C, at a rate of 1°C min^-1^. Equations used for the analysis are described in *Supplementary Information* (Equations S1-S4).

### ATPase activity measurements

ATPase activity of *Af*CopA-NP was determined by measuring the amount of inorganic phosphate (Pi) released from ATP at different reaction times at the temperature stated on each figure. Pi concentration was determined spectrophotometrically according to a modified Malachite Green method ^80,81^. Reaction media contained protein concentrations of 4-6 uM, 25 mM MES pH 5.5 at 70°C, 100 mM NaCl, 30 mM MgCl_2_ and 3 mM ATP unless otherwise indicated. Time course of Pi release was obtained from at least 5 independent reaction tubes after subtraction of ATP hydrolysis in the absence of the protein. Initial rate was calculated as the slope of the corresponding time course with its standard error. Molar ATPase activity was calculated dividing the initial rate by enzyme concentration.

### Model selection

To evaluate the goodness of the model fit we used the Second-Order Akaike Information Criterion (AICc) ^82^ which is defined as:

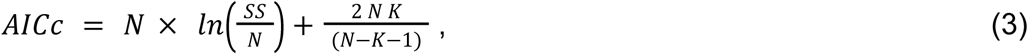

with N = number of data, K = number of parameters plus 1, and SS = sum of weighted square residual errors ^82^. The best model was considered to be the one which gave the lower value of AICc.

### Computational methods

#### Local Frustration analysis

Local Frustration analysis was performed using Frustratometer 2 server ^66^. For a given contact between two residues, its energy is compared to that of an alternative pair of residues in the same location. From the comparison contacts can be classified as highly, minimally or neutrally frustrated. Similarly, frustration for single residues is obtained by comparing its native energy with that obtained from mutating to the other possible amino acid residues. Contacts and residues are classified as highly frustrated when the frustration index is below −1, while they will be minimally frustrated with values above 0.78. Additional details can be found in Ferreiro DU et al. ^83^.

#### AlphaFold2 prediction

AlphaFold2 (AF2) was used to predict the open conformation of the *Af*CopA-NP. We followed the protocol described in Guan et al. ^42^ by first calculating the highly frustrated positions in *Af*CopA-NP (PDB 3A1C A) as described in the previous section. Those positions were masked by introducing a gap in the corresponding column of the AF2 alignment, removing this input information from AlphaFold2 predictions. This protocol has been successfully used to obtain reported conformations for a couple of proteins such as Adenylate kinase, KaiB and RBP ^42^.

#### Structure-based model simulations

We performed dual-basin Cα-Structure-based model simulations (Cα-SBM) using an in-house OpenMM implementation with a force field parameterization described by Onuchic and coworkers ^84^. We used native bonds, angles and dihedrals corresponding to the open structure predicted as described in the previous section. As we evaluated the free energy landscape of the open-closed transition, we modeled the presence of the ligand as the contacts in the closed conformation (PDB ID 3A1C) that are 50% further apart in the open conformation (Q_ligand_ - Figure S14). Thus, native contacts were defined within 2 different basins: the contacts of the open state, with a corresponding energy weight of 1 (ε_nc_=1); and the Q_ligand_, with energy weight variable (ε_Qligand_). The closing reaction was simulated by increasing the ε_Qligand_ favouring the closed conformation as described previously ^84,85^. We biased the simulation from the open to the closed state by employing Umbrella Sampling technique using the difference in the root-mean squared deviation (*RMSD_dif_*) between the reference closed and open states ^42^. The reaction coordinate was calculated using:

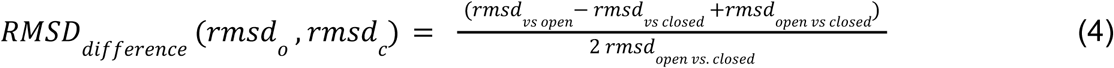

where *rmsd_vs open_*and *rmsd_vs closed_*are the calculated root-mean squared deviation of the current structure with respect to the open and closed states, respectively; *rmsd_open vs closed_* is a constant value representing the root-mean squared deviation between the reference open and closed states. Free energy profiles were obtained by performing several constant temperature simulations and combining them using the Weighted Histogram Analysis Method (WHAM) ^86^. All results in this work are at T = 0.9-0.8 in reduced units (kT = 1). Principal components analysis (PCA) was performed using sklearn python library ^87^ taking two different approaches. On one side, in strain PCA, the covariance matrix was built on the basis of interatomic distance variations among the simulation (d_ij_) between the Cα atoms of two residues, while on the other side, in cracking PCA the covariance matrix was built using a binary definition of contacts and described the fluctuation of the contacts formation; when d_ij_ > 1.2 * native distance the value is 0 while it is 1 otherwise.

## Acknowledgements

This work was funded by *CONICET* (PIP 2022). We thank Dr. José M. Arguello’s (Worcester Polytechnique Institute, Worcester, MA, USA) for sharing the *Af*CopA catalytic unit plasmid. We also thank Dr. Diego U. Ferreiro (Universidad de Buenos Aires) for carefully reading and discussing this manuscript. Additionally, we acknowledge Facundo Balerdi Roman for critical reading. Finally, we manifest our deep concerns about the sudden discontinuation of scientific funding which impedes the development of new projects and terminates the current ones.

