## Supplementary Information for "Cracking Controls ATP Hydrolysis in the catalytic unit of a P-type ATPase"

[e] Consejo Nacional de Investigaciones Científicas y Técnicas (CONICET). Universidad de Buenos Aires. Instituto de Química Biológica de la Facultad de Ciencias Exactas y Naturales (IQUIBICEN). Buenos Aires, Argentina.

Running title: Cracking modulates the ATPase activity of *Af*CopA-NP.

### Supplementary figures

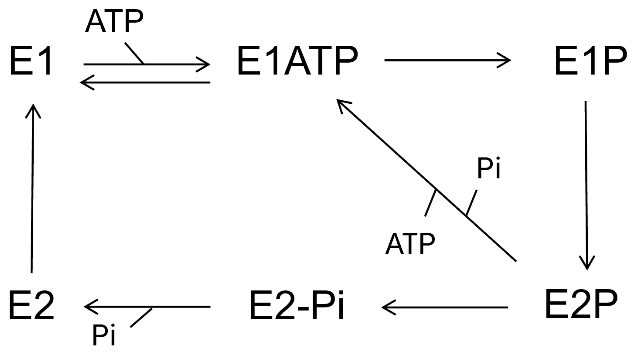


**Figure S1.** Post-Albers reaction cycle scheme. The enzyme can be found in two main conformational ensembles: E1 and E2. Phosphorylated species are denoted as E1P and E2P. ATP can bind to E1 or to E2P as an alternative regulatory mode described for the Sarcoplasmic Reticulum $Ca^{2+}$-ATPase [1].

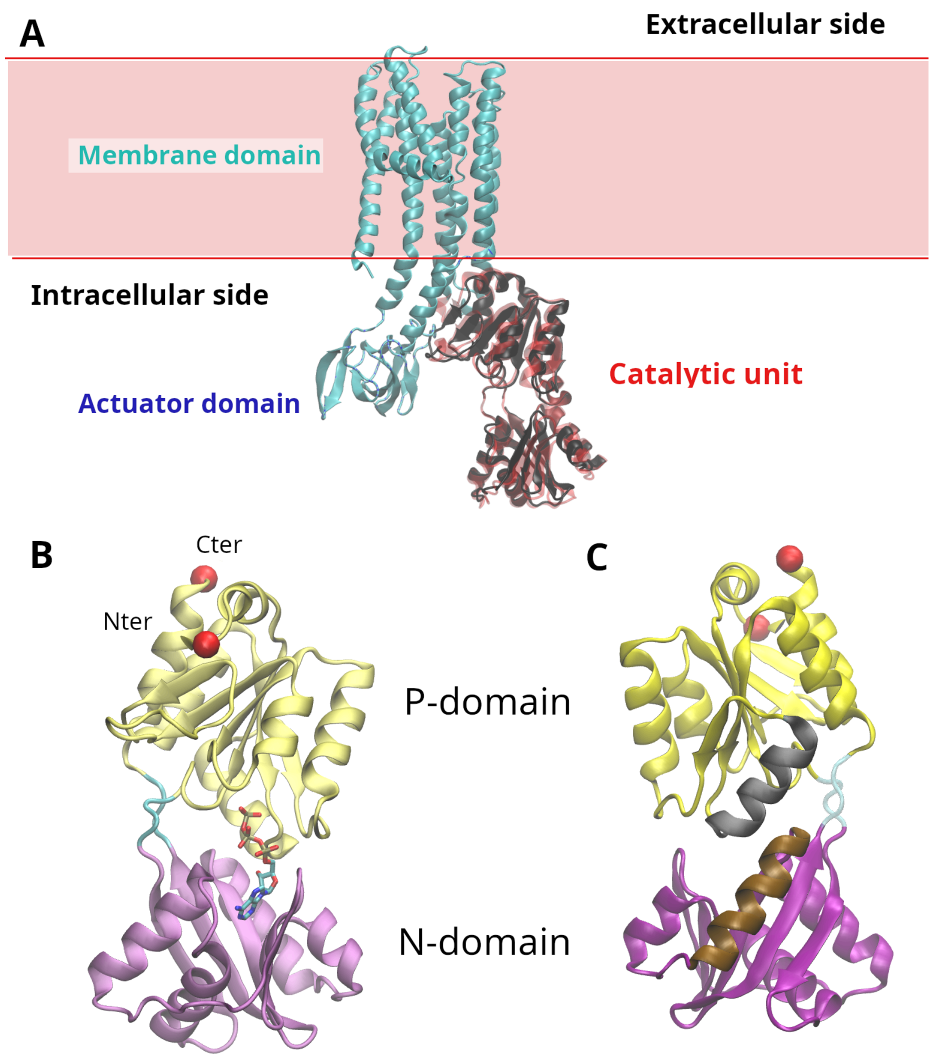


**Figure S2. A**. Structure representation of *Af*CopA (PDB ID 7R0H). Metal ions bind to the transmembrane region while ATP binds to the catalytic unit. After hydrolysis and phosphorylation, the rotation of the Actuator domain concomitantly with dephosphorylation triggers ion transport to the extracellular side. Isolated catalytic unit (PDB ID 3A1C, black) is superimposed on the catalytic unit of *Af*CopA (red). **B**. Structure of the *Af*CopA-NP showing ATP binding site. Red balls represent the N and C termini while in yellow and purple are shown the P and N domains respectively. **C**. Structure of *Af*CopA-NP showing helices M5 and M7 (black and ocre respectively).

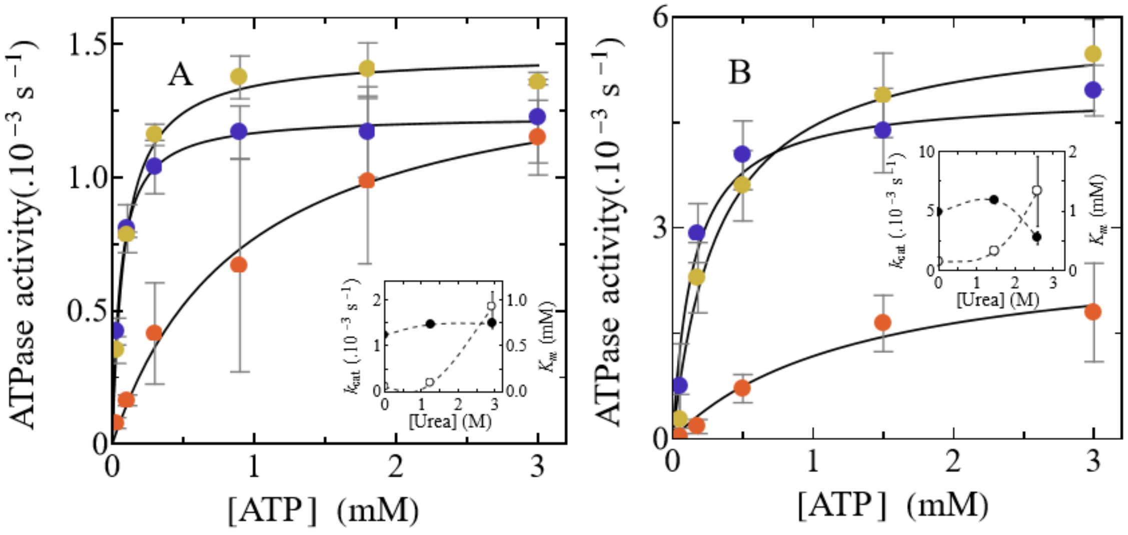


**Figure S3**. ATPase activity dependence on ATP concentration in the presence of urea. **A** ATPase activity determined in the presence of different ATP concentrations at 45 ºC and 0 (●), 1.25 (●) and 3 M urea (●). **B**. ATPase activity determined in the presence of different ATP concentrations at 55 ºC and 0 (●), 1.5 (●) and 2.6 M urea (●). Continuous lines in both panels are the graphical representation of equation 1 fitted to each curve. In all cases, error bars represent the corresponding standard error. Inset in each panel shows the corresponding best fit parameter values for $k_{cat}$ (●) and $K_m$ (○).

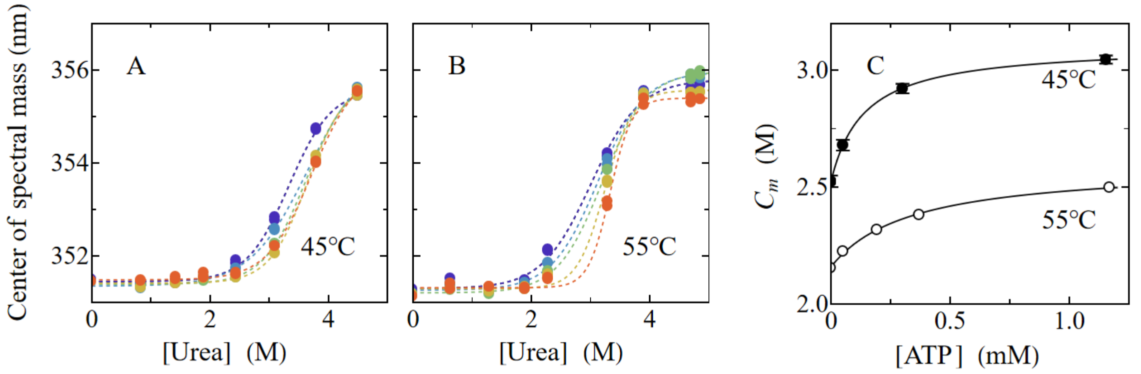


**Figure S4.** Analysis of the effects of urea, ATP and temperature on *Af*CopA-NP intrinsic tryptophan fluorescence. **A**. Trp fluorescence center of spectral mass as a function of urea concentration at 45 and 0 (●), 0.05 (●), 0.3 (●), 1.2 mM ATP (●). **B**. Trp fluorescence center of spectral mass as a function of urea concentration at 55°C and 0 (●), 0.05 (●), 0.2 (●), 0.4 (●) and 1.2 mM ATP (●). Dashed lines constitute a guide for the eye. **C**. Mid-denaturant concentration ($C_m$) values obtained from the dependence of $I_{Trp}$ on urea concentration (Figure 6) as a function of ATP concentration at 45°C (●) or 55°C (o). Continuous lines are the graphical representation of equation $\frac{(a_{inf}-a_0)\,[ATP]}{K_m+[ATP]}+a_0$ fitted to each dataset, the best fit parameter values were $a_0$ = 2.525 ± 0.002 M, $a_{inf}$ = 3.107 ± 0.003 M and $K_m$ = 0.139 ± 0.003 mM at 45°C and $a_0$ = 2.158 ± 0.008 M, $a_{inf}$ = 2.592 ± 0.02 M and $K_m$ = 0.335 ± 0.05 mM at 55°C.

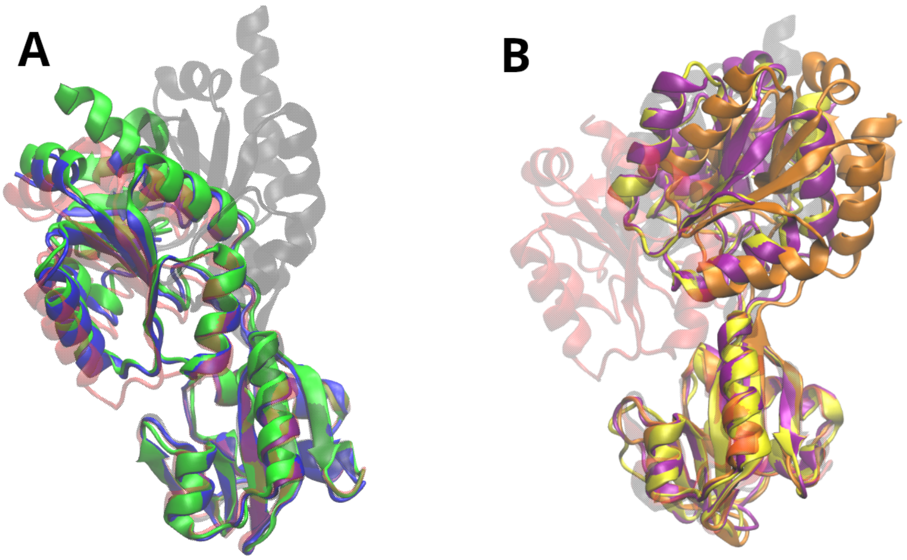


**Figure S5.** Structural alignment of the representative available Cu(I) ATPases catalytic units. **A**. Structures from *Archaeglobus fulgidus* variants, both in the absence (PDB ID 7R0H for the full length protein and 2B8E for the isolated domain, in green and blue, respectively), in the presence of ATP analog (PDB ID 3A1C, in red), and the model of the open conformation predicted using the local frustration/AlphaFold2 protocol (in black). **B**. Structures from *Legionella pneumophila* variants, both in the absence (PDB ID 3RFU in purple), and in E2P and E2-Pi (PDB ID 4BBJ and 4BYG in orange and yellow, respectively). The structures from *Af*CopA-NP in the presence of an ATP analog (PDB ID 3A1C, in red) and the model of the open conformation predicted using the local frustration/AlphaFold2 protocol (in black) are also shown.

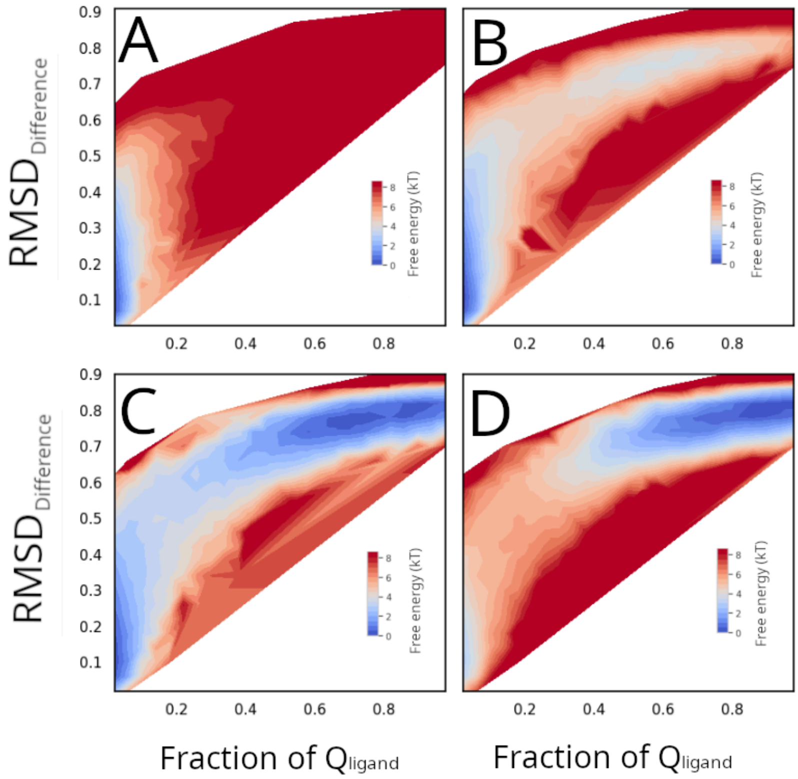


**Figure S6**. Free Energy Landscape of the open-closed transition of *Af*CopA-NP with different energy weights for the $Q_{ligand}$. **A**. $\varepsilon_{Qligand}$ = 0. **B**. $\varepsilon_{Qligand}$ = 0.5. **C**. $\varepsilon_{Qligand}$ = 0.8. **D**. $\varepsilon_{Qligand}$ = 1.

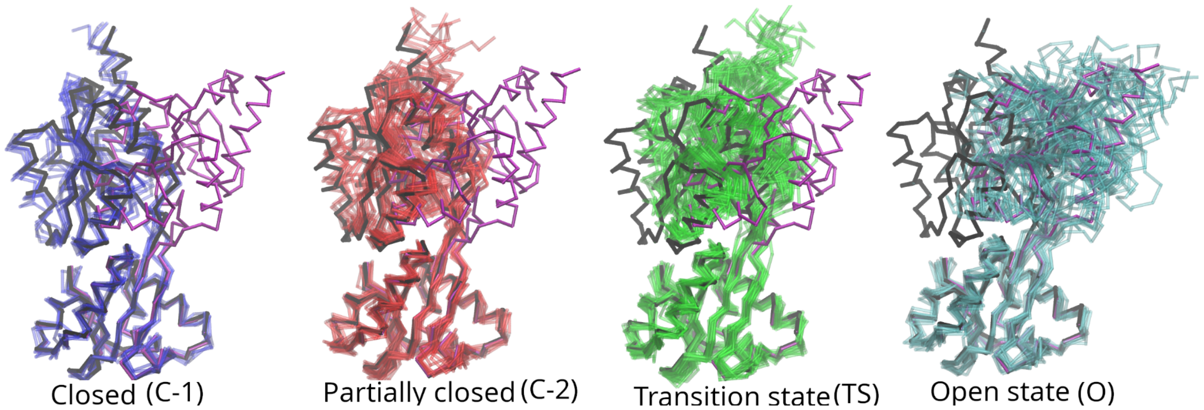


**Figure S7.** Representative structures for the main conformational ensembles of the Free Energy profile shown in Figure 8A. In each panel, black structure is the Cα representation of the PDB ID 3A1C while the purple one corresponds to the AF2 predicted open conformation.

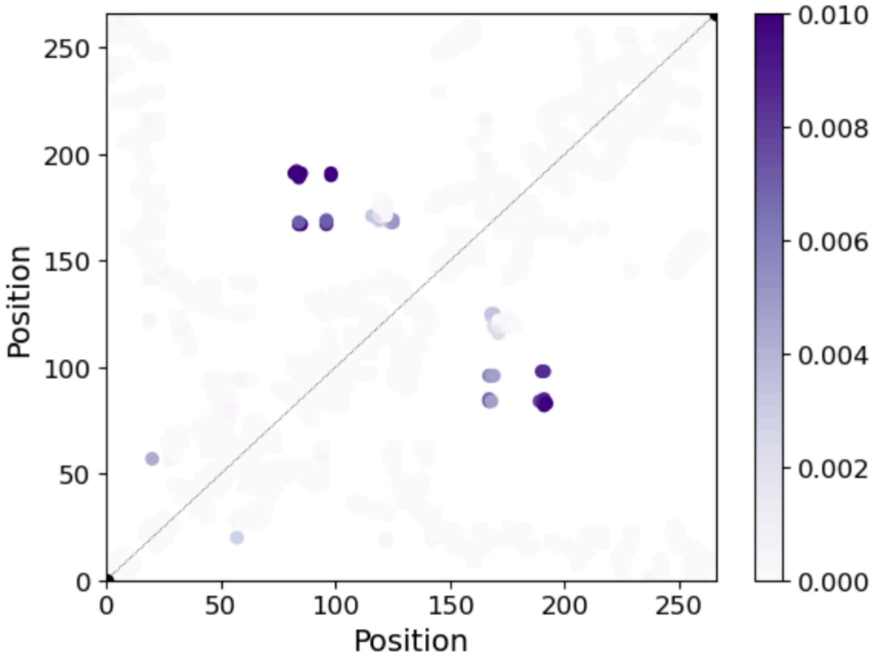


**Figure S8.** Projection of the first principal component of *cracking* and *strain* to the contact map. Contacts above the diagonal represent the cracking PC1 while contacts below the diagonal the strain PC1. Position numbering is according to the *Af*CopA-NP sequence.

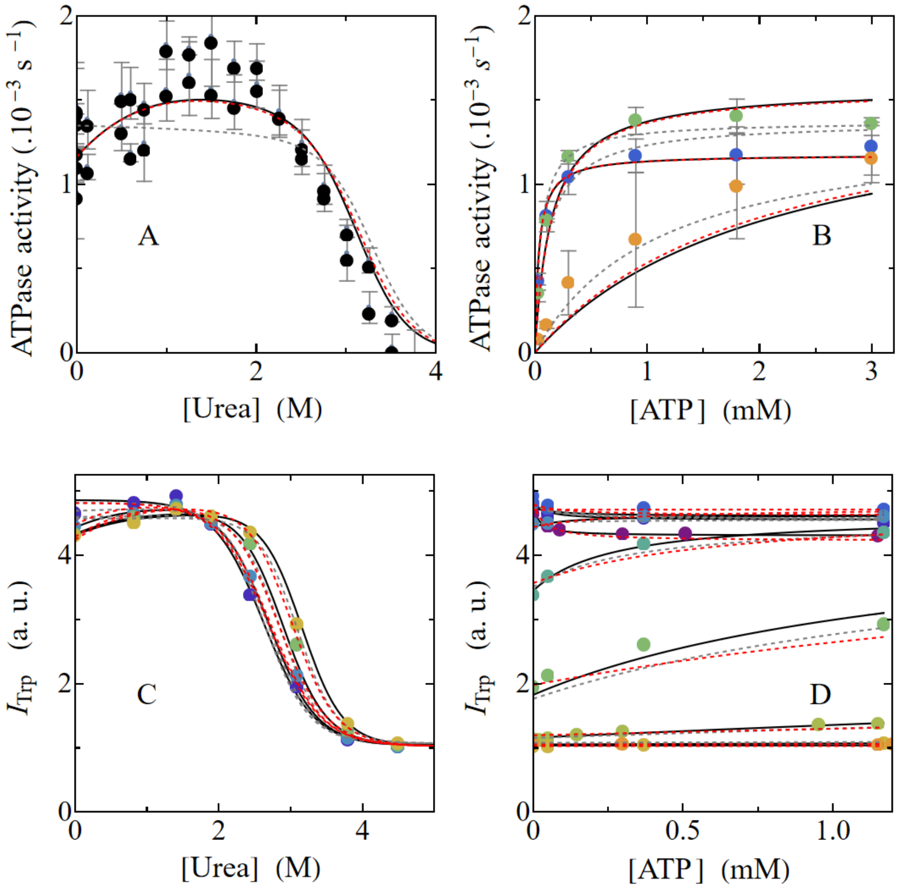


**Figure S9.** Modeling the effect of ATP and urea on ATPase activity and Trp fluorescence at 45°C. **A.** *Af*CopA-NP ATPase activity as a function of urea concentration in the presence of 3 mM ATP. **B.** *Af*CopA-NP ATPase activity as a function of ATP, in the presence of 0 (●), 1.5 (●) or 2.6 M urea (●). **C.** Trp fluorescence intensity at 45 ºC in the presence of different urea concentrations and 0 (●), 0.05 (●), 0.3 (●) and 1.2 mM ATP (●). **D.** The same data plotted in C but as a function of ATP concentration for 0 (●), 0.83 (●), 1.42 (●), 1.9 (●), 2.44 (●), 3.09 (●), 3.8 (●), 4.5 (●) and 5.3 M urea (●). Gray dashed, red dashed and continuous black lines are the graphical representations of the global fit to all the experimental data, of the equations derived from models depicted in Figure 9E (Equations S6 and S7), Figure 9F considering rapid equilibrium condition (Equations S8 and S9) or Figure 9F considering ATP binding out-of the rapid equilibrium condition (Equations S8 and S10), respectively.

Best fit parameter values are:

| $K_1$ | $k_2$ ($s^{-1}$ $mM^{-1}$) | $k_{-2}$ ($s^{-1}$) | $K_3$ | $k_{AATP}$ ($s^{-1}$) |
|---|---|---|---|---|
| 0.0004 ± 0.0003 | 10 ± 3 | 0.16 ± 0.04 | 0.5 ± 0.1 | (1.7 ± 0.1) .10$^{-3}$ |

| $m_1$ (kcal $mol^{-1}$ $M^{-1}$) | $m_2$ (kcal $mol^{-1}$ $M^{-1}$) | $m_{-2}$ (kcal $mol^{-1}$ $M^{-1}$) | $m_3$ (kcal $mol^{-1}$ $M^{-1}$) |
|---|---|---|---|
| -1.8 ± 0.1 | 0.6 ± 0.06 | 0 | 0.9 ± 0.4 |

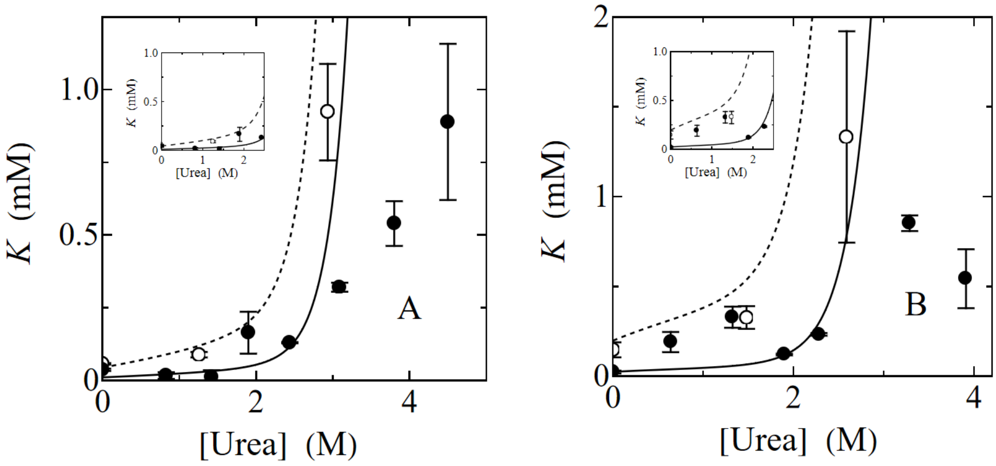


**Figure S10.** Model prediction of urea dependence of ATPase activity $K_m$ and Trp fluorescence $K_{ATP}$. **A, B**. $K_m$ values (o) were obtained from the dependence of the ATPase activity on substrate concentration (Figure S3). Values of the dissociation constant $K_{ATP}$ (●) were obtained from the best fit of equation S5 to tryptophan fluorescence (FIgure 6) as a function of urea concentration at 45°and 55°C respectively. Dashed and continuous lines correspond to the predictions of $K_m$ and $K_{ATP}$ from the model of Figure 9F considering the ATP binding step out-of the rapid equilibrium with the parameter values shown in Tables S4 and S5.

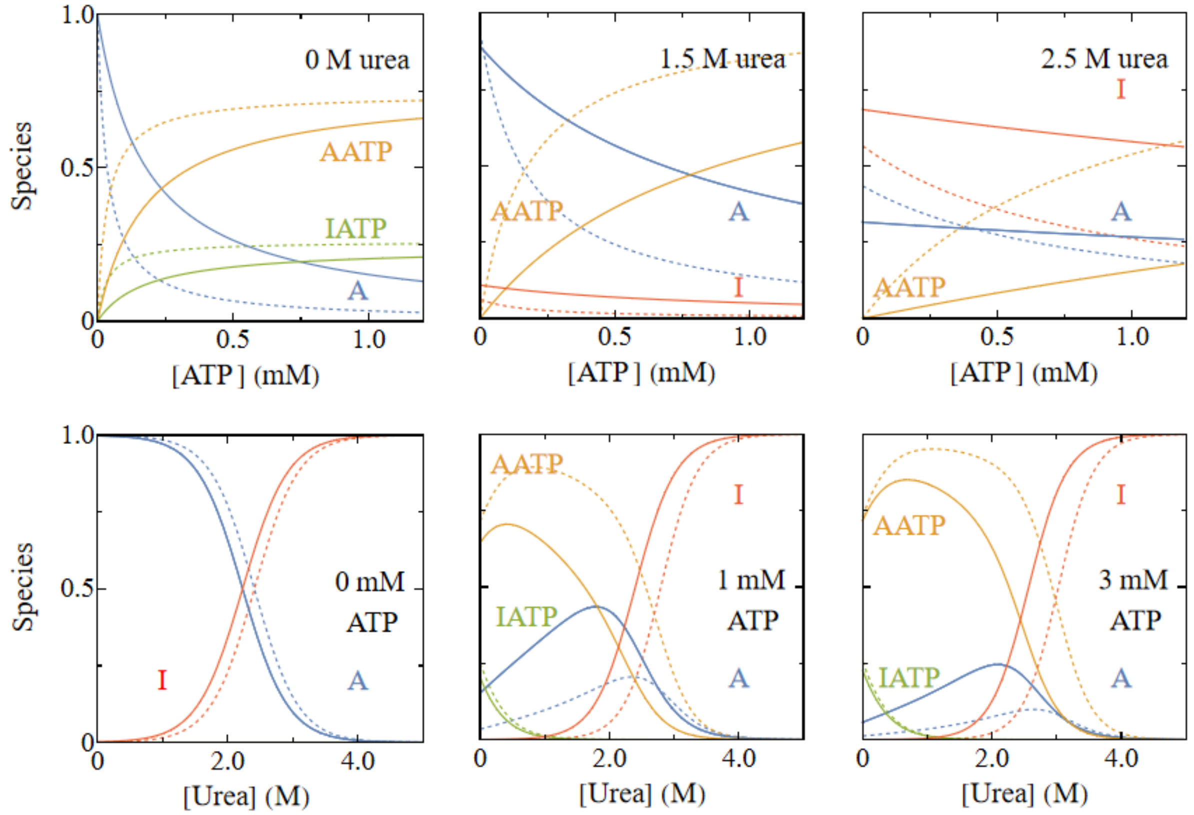


**Figure S11**. Modelling the effect of ATP, urea and temperature on the enzyme species distribution. Simulation of the molar fraction of the species depicted in the model from Figure 9F (A, I, AATP and IATP) as a function of ATP or urea concentration at 55°C (continuous lines) and 45°C (dashed lines). Simulations were performed using the model derived equations for each species and the best fit parameter values from Table S6 or Figure S9, respectively.

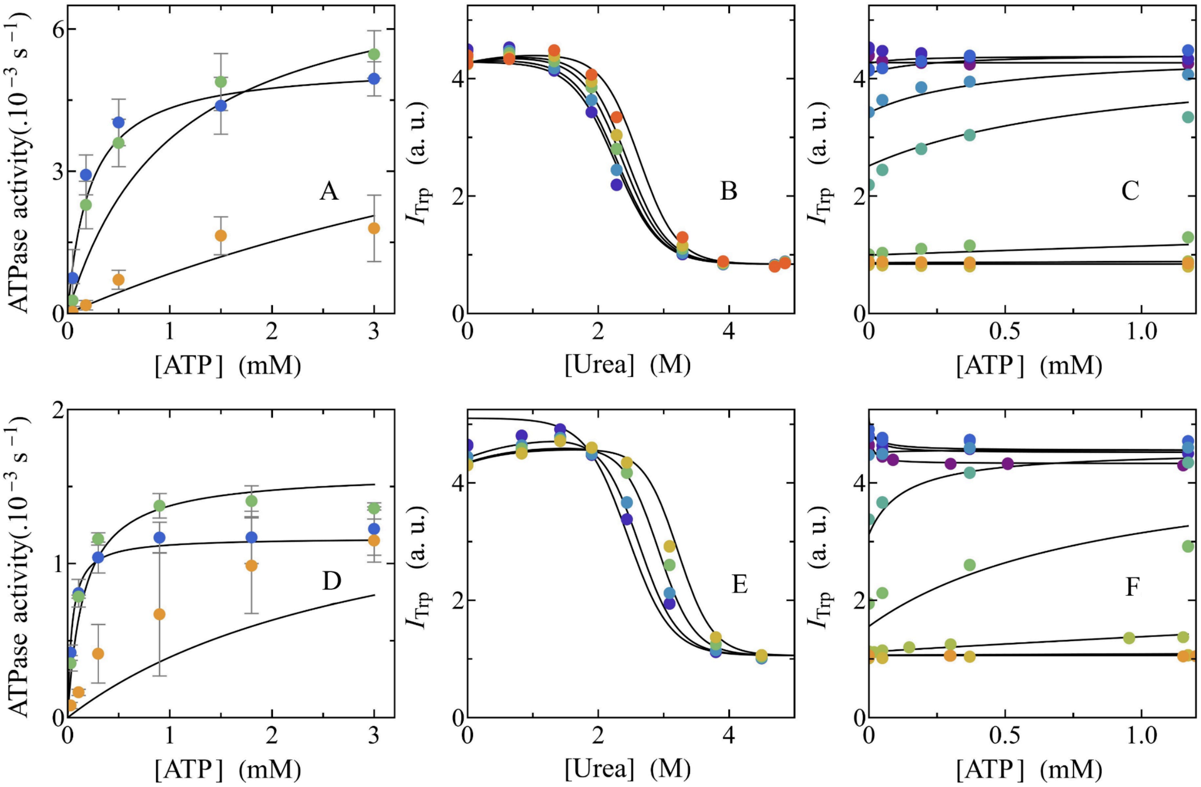


**Figure S12.** Modelling the effect of urea, ATP and temperature on intrinsic tryptophan fluorescence and ATPase activity. **A** and **D**. ATPase activity determined in the presence of different ATP concentrations at 55 ºC and 0 (●), 1.5 (●) and 2.6 M urea (●), or at 45 ºC and 0 (●), 1.25 (●) and 3 M urea, respectively. **B** and **E**. tryptophan fluorescence ($I_{Trp}$) at different urea concentrations at 55°C and 0 (●), 0.05 (●), 0.2 (●), 0.4 (●) and 1.2 mM ATP (●) or at 45°C and 0 (●), 0.05 (●), 0.3 (●) and 1.2 mM ATP (●), respectively. **C** and **F**. Tryptophan fluorescence ($I_{Trp}$) at different ATP concentrations at 55°C and 0 (●), 0.6 (●), 1.3 (●), 1.9 (●), 2.3 (●), 3.3 (●), 4.0 (●) and 4.8 M urea (●) or at 45°C and 0 (●), 0.83 (●), 1.42 (●), 1.9 (●), 2.44 (●), 3.09 (●), 3.8 (●), 4.5 (●) and 5.3 M urea (●), respectively. Continuous lines are the graphical representation of the equations derived from the model shown in Figure 9F (Equations S11 and S12) fitted to both spectroscopic and molar ATPase activity data at all temperatures, ATP and urea concentrations assayed. Best fit parameter values are shown in Table S3.

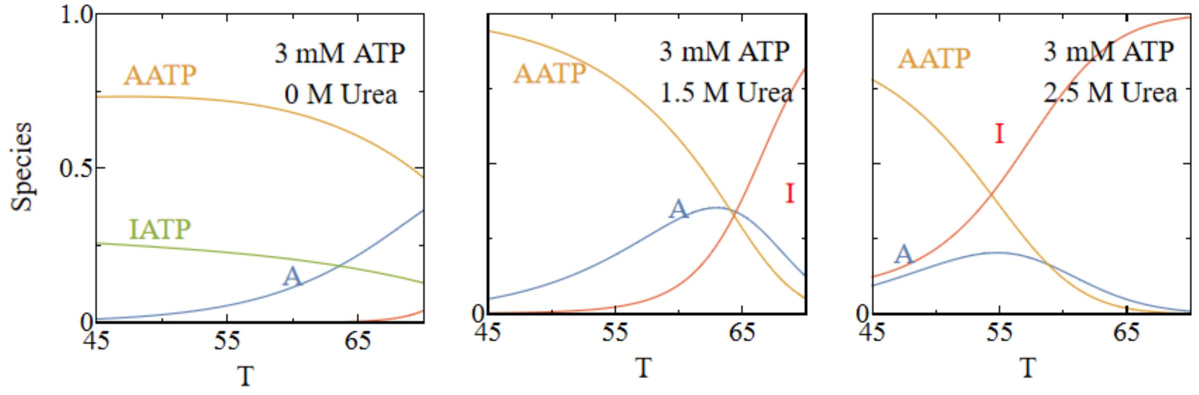


**Figure S13**. Modelling the effect of ATP, urea and temperature on the enzyme species. Simulation of the molar fraction of the species depicted in the model from Figure 9F (A, I, AATP and IATP) as a function of temperature at the ATP and urea concentrations stated in each figure.

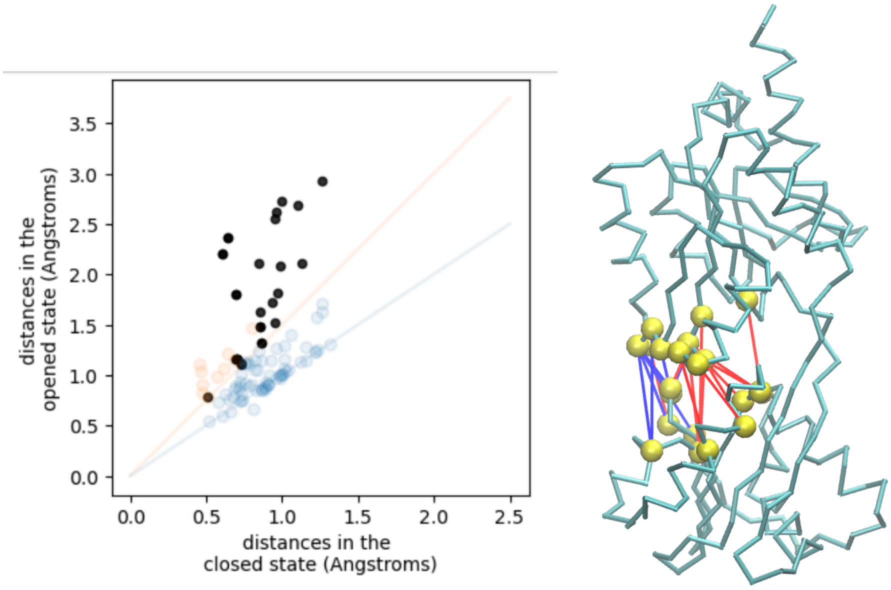


**Figure S14.** $Q_{ligand}$ definition. **A**. Each point represents the distance between the Cα of the residues involved in the contact, in the closed and in the open conformation. The blue line is a straight line with slope = 1 and intercept = 0, while the orange line is a straight line with slope = 1.5 and intercept = 0. Blue dots represent the contacts unique to the closed conformation whose distance increases less than 1.5 times in the open conformation. Orange dots are the contacts unique in the closed conformation whose distance increase more than 1.5 times in the open conformation but are within N or P domains. Black dots are contacts unique in the closed conformation whose distance increases more than 1.5 times in the open state and are between both domains (each one of these dots is considered as a $Q_{ligand}$). **B**. Representation of ligand mediated contacts (provided as a Supplementary list - List S1). In yellow spheres are the alpha carbons of residues involved, red and blue lines represent the contacts involved in C1 ensemble, and in blue are the C1 ensemble contacts disrupted in C2.

#### TABLES

**Table S1.** AICc values comparison for the fitting of 2-states (Equation S1) or 3-states (Equation S2) models to the experimental data shown in Figure 2.

| *2-states model* | *3-states model* |
|---|---|
| -344 | -376 |

**Table S2.** Best fit parameter values of Equation S2 fitted to the values of $I_{Trp}$ and $[\theta]_{220nm}$ as a function of temperature (Figure 2).

| $\Delta H_{AI}$ (kcal mol$^{-1}$) | $\Delta Cp_{AI}$ (kcal mol$^{-1}$K$^{-1}$) | $Tm_{AI}$ (°C) | $\Delta H_{IU}$ (kcal mol$^{-1}$) | $\Delta Cp_{IU}$ (kcal mol$^{-1}$K$^{-1}$) | $Tm_{IU}$ (°C) |
|---|---|---|---|---|---|
| 77 ± 3 | 2.0 ± 0.1 | 72.0 ± 0.2 | 86 ± 8 | 2.3 ** | 86.7 ± 0.3 |
| $Sa_f$ (a.u.) | $Si_f$ (a.u.) | $Su_f$ (a.u.) | $ma_f$ (a.u.) | $mi_f$ (a.u.) | $mu_f$ (a.u.) |
| 6.40 ± 0.09 | 0.149 ± 0.008 | 0.149 ± 0.008* | -0.8 ± 0.5 | 0 * | 0 * |
| $Sa_{cd}$ (a.u.) | $Si_{cd}$ (a.u.) | $Su_{cd}$ (a.u.) | $ma_{cd}$ (a.u.) | $mi_{cd}$ (a.u.) | $mu_{cd}$ (a.u.) |
| - 6.7 ± 0.1 | -2.78 ± 0.03 | -1.94 ± 0.02 | 0.0094 ± 0.0004 | 0 * | 0 * |

*$Su_f$ was fixed to be equal to $Su_i$, and $mi_f$, $mu_f$, $mi_{cd}$ and $mu_{cd}$ were fixed equal to 0 according to Akaike Information Criterion [2] .

**$\Delta Cp_2$ was fixed so that $\Delta Cp_{IA}$ + $\Delta Cp_{UI}$ equals the theoretical $\Delta Cp$ for AfCopA-NP (4.3 *kcal mol$^{-1}$K$^{-1}$*) according to Myers et al. [3].

**Table S3.** Best fit parameter values of the equations derived from the model shown in Figure 9 F without rapid equilibrium assumption and including temperature dependences (Equations S11 and S12) fitted to the experimental data shown in Figures 3 and S12.

| $K_1$ | $k_2$ ($s^{-1}$ $mM^{-1}$) | $k_{-2}$ ($s^{-1}$) | $K_3$ | $k_{AATP}$ ($s^{-1}$) | |
|---|---|---|---|---|---|
| 0.0011 ± 0.0005 | 7 ± 1 | 0.5 ± 0.2 | 0.44 ± 0.09 | (7.6 ± 0.8).$10^{-3}$ | |
| $m_1$ (kcal $mol^{-1}$ $M^{-1}$) | $m_2$ (kcal $mol^{-1}$ $M^{-1}$) | $m_{-2}$ (kcal $mol^{-1}$ $M^{-1}$) | $m_3$ (kcal $mol^{-1}$ $M^{-1}$) | | |
| -1.9 ± 0.1 | 0.49 ± 0.04 | 0 | 1.3 ± 0.4 | | |
| $\Delta H_{K1}$ (kcal $mol^{-1}$) | $\Delta C_p$ (kcal $mol^{-1}K^{-1}$) | $E_{Ak2}$ (kcal $mol^{-1}$) | $E_{Ak-2}$ (kcal $mol^{-1}$) | $\Delta H_{K3}$ (kcal $mol^{-1}$) | $E_{AkAATP}$ (kcal $mol^{-1}$) |
| 36 ± 3 | 1.9 ± 0.2 | 0 ± 4 | 43 ± 9 | 2 ± 4 | 31 ± 1 |

**Table S4.** Best fit parameter values of Equation S1 fitted to $I_{Trp}$ as a function of temperature and urea concentration shown in Figure 5.

| $\Delta H_1$ (kcal $mol^{-1}$) | $\Delta Cp_1$ (kcal $mol^{-1}K^{-1}$) | $Tm_1$ (°C) | | | |
|---|---|---|---|---|---|
| 88 ± 6 | 2.4 ± 0.2 | 71.2 ± 0.3 | | | |
| $Sn_f$ (a.u.) | $Su_f$ (a.u.) | $mn_f$ (a.u.) | $mu_f$ (a.u.) | $In_f$ (a.u.) | $Iu_f$ (a.u.) |
| 6.7 ± 1 | 0.6 ± 0.02 | -0.48 ± 0.02 | 0 * | 0.8 ± 0.5 | 0 * |

* $mu_f$ and $Iu_f$ were fixed equal to 0 according to Akaike Information Criterion [2] .

**Table S5.** AICc values for comparison of fittings of the model derived equations from Figure 9E (Equations S6 and S7), Figure 9F under rapid-equilibrium assumption (Equations S8 and S9) or Figure 9F without the rapid-equilibrium assumption (Equations S8 and S10) to the ATPase activity and $I_{Trp}$ results at 45°C (Figure S10) or at 55°C (Figure 9).

| *Temperature* | *Scheme 2* *Rapid-equilibrium* | *Scheme 3* *Rapid-equilibrium* | *Scheme 3* *Out of Rapid-equilibrium* |
|---|---|---|---|
| 45 | -166 | -198 | -210 |
| 55 | -388 | -393 | -439 |

**Table S6.** Best fit parameter values of the equations derived from the best model (Figure 9F without rapid equilibrium, Equations S8 and S10) fitted to the experimental data determined at 55°C shown in Figure 9.

| $K_1$ | $k_2$ ($s^{-1}$ $mM^{-1}$) | $k_{-2}$ ($s^{-1}$) | $K_3$ | $k_{AATP}$ ($s^{-1}$) |
|---|---|---|---|---|
| 0.0016 ± 0.0006 | 7.5 ± 0.9 | 0.4 ± 0.1 | 0.3 ± 0.1 | (7.1 ± 0.7) .10$^{-3}$ |

| $m_1$ (kcal $mol^{-1}$ $M^{-1}$) | $m_2$ (kcal $mol^{-1}$ $M^{-1}$) | $m_{-2}$ (kcal $mol^{-1}$ $M^{-1}$) | $m_3$ (kcal $mol^{-1}$ $M^{-1}$) |
|---|---|---|---|
| -1.9 ± 0.1 | 0.41 ± 0.04 | 0 | 0.8 ± 0.5 |

#### EQUATIONS

**Two-states unfolding transitions (A⇌U)**:

$$S = \frac{(Sa + ma\ D + Ia\ T) + (Su + mu\ D + Iu\ T)\ K_{eq\ U-A}}{1 + K_{eq\ U-A}} \qquad \text{(S1)}$$

where $S_a$, and $S_u$ are the folded and unfolded signals at equilibrium, respectively. $m_a$, $I_a$, $m_u$ and $I_u$ are the signal dependencies of the folded, intermediate and unfolded conformations with urea and temperature, respectively, while D is the urea concentration. The equilibrium constant was defined as $K_{eq\ U\text{-}A}$ = U/A,

**Three-states unfolding transitions**:

$$S = \frac{(Sa + ma\ D + Ia\ T) + (Si + mi\ D + Ii\ T)\ K_{eq\ I-A} + (Su + mu\ D + Iu\ T) K_{eq\ U-I}\ K_{eq\ I-A}}{1 + K_{eq\ I-A} + K_{eq\ U-I}\ K_{eq\ I-A}} \qquad \text{(S2)}$$

where $S_a$, $f_a$, $S_I$, $f_I$, $S_u$ and $f_u$ are the folded, intermediate and unfolded signals and fractions at equilibrium, respectively. $m_{na}$, $I_a$, $m_i$, $I_I$, $m_u$ and $I_u$ are the signal dependencies of the folded, intermediate and unfolded conformations with urea concentration and temperature, respectively, while D is the urea concentration. Equilibrium constants were defined as $K_{eq\ U\text{-}I}$ = U/I and $K_{eq\ I\text{-}A}$ = I/A.

In both cases,

$$K_{eq} = K^0_{eq}\ e^{-m_{eq} D} \qquad \text{(S3)}$$

where $K^0_{eq}$ corresponds to the equilibrium constant in the absence of urea and *m* is the slope of the dependence of $Ln\ (K_{eq})$ with the denaturant concentration.

and:

$$K^0_{eq} = e^{\frac{-\left(\Delta H_m \left(1 - \frac{T}{T_m}\right) + \Delta Cp \left(T - T_m - T\ Ln\left(\frac{T}{T_m}\right)\right)\right)}{R\,T}} \qquad \text{(S4)}$$

where T corresponds to the temperature, ΔH to the enthalpy change, $T_m$ the temperature where the equilibrium constant equals 1, ΔCp to the heat capacity change and R is the gas constant (0.001987 kcal $K^{-1}$ $M^{-1}$ ).

**ATP binding**

$$S = S_E \frac{1}{2}\left(- ATP_t + E_t - K_{ATP} + \sqrt{- 4\, ATP_t E_t + (- ATP_t - E_t - K_{ATP})^2}\right)$$
$$+ S_{EATP} \frac{1}{2}\left(ATP_t + E_t + K_{ATP} - \sqrt{- 4\, ATP_t E_t + (- ATP_t - E_t - K_{ATP})^2}\right) \qquad \text{(S5)}$$

$E_T$ and $ATP_T$ correspond to the total enzyme and ATP concentrations. E and E.ATP correspond to the free and ATP bound enzyme species, respectively. $K_{ATP}$ is the dissociation constant of the E.ATP complex. $S_E$ and $S_{EATP}$ correspond to the fluorescence intensity of E and E.ATP respectively. This equation was obtained considering that $E_t$ = E + E.ATP and $ATP_t$ = E.ATP + ATP.

**Model from Figure 9E**

**Under rapid equilibrium assumption**:

$$S = Et\,\frac{S_A K_{1\,ref}\, e^{\frac{-m1\,D}{R\,T}} + S_{AATP} ATP + S_I\, K_{I-A\,ref} K_{1\,ref}\, e^{\frac{-(m_{I-A}+m1)\,D}{R\,T}}}{K_{1\,ref}\, e^{\frac{-m1\,D}{R\,T}} + ATP + K_{I-A\,ref} K_{1\,ref}\, e^{\frac{-(m_{I-A}+m1)\,D}{R\,T}}} \qquad \text{(S6)}$$

$$ATPase\ activity = \frac{k_{AATP} ATP}{K_{1\,ref}\, e^{\frac{-m1\,D}{R\,T}} + ATP + K_{I-A\,ref} K_{1\,ref}\, e^{\frac{-(m_{I-A}+m1)\,D}{R\,T}}} \qquad \text{(S7)}$$

$E_T$, ATP and D correspond to the total enzyme, ATP and urea concentrations. Equilibrium dissociation constants and rate coefficients are defined in terms of the reactions depicted in the model from Figure 9E. Equilibrium constant dependence on urea concentration is given by $K_0 \cdot e^{-m\,D/R\,T}$. Equation was obtained considering that all enzyme species are in rapid equilibrium. $S_A$, $S_{AATP}$ and $S_I$ correspond to the fluorescence intensity of the enzyme species A, A.ATP and I respectively.

**Model from Figure 9F**

**Under rapid equilibrium assumption:**

$$S = Et \frac{S_A\ K_{2\,ref}\, e^{\frac{-m2\ D}{R\ T}} + S_{AATP}ATP + S_{IATP}ATP\ K_{3\,ref}\, e^{\frac{-m3\ D}{R\ T}} + S_I\, K_{1\,ref}\, K_{2\,ref}\ e^{\frac{-(m1+m2)\ D}{R\ T}}}{K_{2\,ref}\, e^{\frac{-m2\ D}{R\ T}} + ATP + ATP\ K_{3\,ref}\, e^{\frac{-m3\ D}{R\ T}} + K_{1\,ref}\, K_{2\,ref}\ e^{\frac{-(m1+m2)\ D}{R\ T}}} \qquad \text{(S8)}$$

$$ATPase\ activity = \frac{k_{AATP}\ ATP\ K_{2\,ref}\, e^{\frac{-m2\ D}{R\ T}}}{K_{2\,ref}\, e^{\frac{-m2\ D}{R\ T}} + ATP\ + ATP\ K_{3\,ref}\, e^{\frac{-m3\ D}{R\ T}} + K_{1\,ref}\, K_{2\,ref}\ e^{\frac{-(m1+m2)\ D}{R\ T}}} \qquad \text{(S9)}$$

$E_T$, ATP and D correspond to the total enzyme, ATP and urea concentrations. The equilibrium dissociation constants and rate coefficients are defined in terms of the reactions depicted in the model from Figure 9F. The equilibrium constant dependence on urea concentration is given by $K_0 \cdot e^{-m\,D\,/\,R\,T}$ . Equations were obtained considering that all enzyme species are in rapid equilibrium. $S_A$ , $S_{AATP}$ , $S_{IATP}$ and $S_I$ correspond to the fluorescence intensity of the enzyme species A, A.ATP, I.ATP and I respectively.

**Without rapid equilibrium assumption:**

Defining $K_2 = k_{-2} / k_2$ , ATPase activity is described by:

$$ATPase\ activity = \frac{k_{AATP}\, ATP}{ATP + ATP\ K_{3\,ref}\, e^{\frac{-m3\,D}{R\,T}} + \frac{k_{AATP}}{k_{2\,ref}}\, e^{\frac{m2\,D}{R\,T}} + k_{AATP}\, \frac{K_{1\,ref}}{k_{2\,ref}}\, e^{\frac{(-m1+m2)\,D}{R\,T}} + \frac{k_{-2\,ref}}{k_{2\,ref}}\, e^{\frac{(m2-mi2)\,D}{R\,T}} + \frac{K_{1\,ref}\, k_{-2\,ref}}{k_{2\,ref}}\, e^{\frac{(-m1+m2-mi2)\,D}{R\,T}}}$$

(S10)

$E_T$, ATP and D correspond to the total enzyme, ATP and urea concentrations. The equilibrium dissociation constants and rate coefficients are defined in terms of the reactions depicted in the model from Figure 9F. The equilibrium constant dependence on urea concentration is given by $K_0 \cdot e^{-m\,D\,/\,R\,T}$ .

**Incorporating temperature dependence:**

$$S = Et\,\frac{S_A + S_{AATP} ATP \frac{k_{-2\,ref}}{k_{2\,ref}}\, e^{\frac{-(m2-m_{-2})\,D}{R\,T}}\, e^{\frac{-(Ea2-Ea_{-2})\,(Tref-T)}{R\,T\,Tref}} + S_{IATP}\, ATP\, K_{3\,ref} \frac{k_{-2\,ref}}{k_{2\,ref}}\, e^{\frac{-(m2-m_{-2}+m3)\,D}{R\,T}}\, e^{\frac{-(Ea2-Ea_{-2})\,(Tref-T)}{R\,T\,Tref}}\, e^{\frac{-\Delta H3\ (1/T-1/Tref)}{R}} + S_I\, K_{1\,ref}\, e^{\frac{-m1\,D}{R\,T}}\, e^{\frac{-\Delta H1+\Delta Cp\,(T-Tref)\ (1/T-1/Tref)}{R}}}{1 + ATP \frac{k_{-2\,ref}}{k_{2\,ref}}\, e^{\frac{-(m2-m_{-2})\,D}{R\,T}}\, e^{\frac{-(Ea2-Ea_{-2})\,(Tref-T)}{R\,T\,Tref}} + ATP\, K_{3\,ref} \frac{k_{-2\,ref}}{k_{2\,ref}}\, e^{\frac{-(m2-m_{-2}+m3)\,D}{R\,T}}\, e^{\frac{-(Ea2-Ea_{-2})\,(Tref-T)}{R\,T\,Tref}}\, e^{\frac{-\Delta H3\ (1/T-1/Tref)}{R}} + K_{1\,ref}\, e^{\frac{-m1\,D}{R\,T}}\, e^{\frac{-\Delta H1+\Delta Cp\,(T-Tref)\ (1/T-1/Tref)}{R}}}$$

(S11)

$ATPase\ activity = Et\,.$

$$\frac{\left(Et\, k_{AATP\,ref}\ ATP \frac{k_{2\,ref}}{k_{-2\,ref}}\, e^{\frac{-(m2-m_{-2})\,D}{R\,T}}\, e^{\frac{-(Ea2-Ea_{-2}+Ea_{kAATP})\,(Tref-T)}{R\,T\,Tref}}\right)}{1 + ATP \frac{k_{2\,ref}}{k_{-2\,ref}}\, e^{\frac{-(m2-m_{-2})\,D}{R\,T}}\, e^{\frac{-(Ea2-Ea_{-2})\,(Tref-T)}{R\,T\,Tref}} + ATP\, K_{3\,ref} \frac{k_{2\,ref}}{k_{-2\,ref}}\, e^{\frac{-(m2-m_{-2}+m3)\,D}{R\,T}}\, e^{\frac{-(Ea2-Ea_{-2})\,(Tref-T)}{R\,T\,Tref}}\, e^{\frac{-\Delta H3\ (1/T-1/Tref)}{R}} + \frac{K_{1\,ref}}{k_{-2\,ref}} k_{AATP\,ref}\, e^{\frac{-(m1+m_2)\,D}{R\,T}}\, e^{\frac{-(Ea_{kAATP}-Ea_{-2})\,(Tref-T)}{R\,T\,Tref}}\, e^{\frac{-\Delta H1+\Delta Cp\,(T-Tref)\ (1/T-1/Tref)}{R}} + K_{1\,ref}\, e^{\frac{-m1\,D}{R\,T}}\, e^{\frac{-\Delta H1+\Delta Cp\,(T-Tref)\ (1/T-1/Tref)}{R}})}$$

(S12)

Temperature dependence was also incorporated in the equations derived from the reaction model shown in Figure 9F without including rapid equilibrium assumption. The dependence of the equilibrium constant on urea concentration is as previously stated. The dependence of the equilibrium constants K1 and $K_3$ on temperature is given by Van’t Hoff equation $K_{ref}\ e^{-\Delta H/R\ (1/T\ -\ 1/Tref)}$. In the case of $K_1$ a dependence of ΔH with temperature $\Delta H = \Delta H_{ref} + \Delta Cp\ (T - T_{ref})$ is incorporated. Reaction rates dependence on temperature is given by an Arrhenius dependence $k_{ref}\ e^{-Ea\ (Tref-T)\ /\ (R\ T\ Tref)}$.

$$\alpha_{contact,i} = \frac{q_{ij(partially\ closed\ \mathrm{or}\ transition\ state)} - q_{ij(open\ state)}}{q_{ij(closed\ state)} - q_{ij(open\ state)}} \quad \text{(S13)}$$

$q_{i,j}$ represents the frequency of the contact between atoms i and j in the partially closed or transition state, with respect to the open and closed states. The value goes from 1, when the contact is formed in the transition state in the same frequency as in the closed state, to 0, when the contact is not formed in that state.

**List S1** Complete lists of contact sets and distances (Å): (QLigand, σligand), (Qopen, σopen) & (Qclosed, σclosed)

QLigand, N=22 (Residue i, Residue j, σij )

[(20, 57, 8.65), (83, 191, 6.39), (83, 192, 9.57), (84, 191, 6.03), (84, 167, 6.96), (96, 167, 11.32), (98, 190, 11.01), (98, 191, 12.63), (116, 171, 7.28), (119, 172, 5.08), (119, 169, 7.26), (124, 168, 7.03), (124, 169, 6.98), (125, 169, 8.57), (82, 191, 9.94), (84, 168, 8.54), (84, 189, 8.41), (85, 167, 9.89), (85, 191, 9.55), (96, 168, 9.73), (96, 169, 9.31), (125, 168, 9.52)]

Qopen, N=857 (Residue i, Residue j, σij )

[(0, 242, 4.31), (0, 243, 6.13), (0, 5, 7.06), (0, 241, 6.05), (0, 6, 7.89), (0, 255, 8.67), (0, 258, 11.21), (0, 9, 10.54), (0, 259, 10.6), (1, 5, 7.6), (1, 240, 8.01), (1, 241, 5.46), (1, 242, 6.05), (1, 243, 7.65), (1, 238, 9.29), (1, 237, 9.87), (2, 241, 6.13), (2, 6, 7.65), (3, 7, 7.48), (4, 8, 6.36), (4, 9, 8.59), (5, 225, 6.63), (5, 241, 6.08), (5, 242, 7.29), (5, 9, 5.88), (5, 259, 8.53), (6, 259, 5.04), (6, 262, 7.92), (6, 263, 8.04), (6, 10, 6.63), (6, 255, 8.25), (6, 258, 8.15), (7, 263, 6.99), (7, 11, 7.11), (7, 259, 6.73), (8, 206, 8.32), (8, 225, 6.63), (8, 12, 6.55), (8, 224, 6.89), (9, 225, 6.52), (9, 256, 5.47), (9, 242, 9.49), (9, 255, 7.65), (9, 259, 5.98), (9, 160, 11.26), (9, 260, 7.69), (10, 259, 5.35), (10, 256, 6.17), (10, 260, 5.0), (10, 263, 7.28), (11, 206, 6.65), (12, 260, 10.02), (12, 206, 4.92), (12, 160, 7.26), (12, 256, 8.0), (12, 207, 5.93), (12, 208, 6.24), (12, 224, 7.92), (12, 225, 7.83), (13, 206, 5.2), (13, 208, 7.98), (13, 159, 7.52), (13, 160, 4.94), (13, 161, 6.19), (13, 205, 5.7), (13, 204, 9.2), (14, 205, 6.0), (14, 206, 6.09), (14, 207, 4.97), (14, 160, 5.37), (14, 161, 4.49), (14, 184, 9.73), (14, 208, 6.2), (15, 160, 6.92), (15, 161, 6.21), (15, 162, 5.01), (15, 207, 5.9), (15, 208, 4.72), (15, 155, 8.02), (15, 163, 6.19), (15, 210, 7.28), (15, 227, 9.72), (15, 252, 9.23), (15, 256, 10.26), (16, 207, 7.08), (16, 208, 6.17), (16, 209, 4.9), (16, 210, 6.09), (16, 163, 4.6), (16, 165, 6.75), (16, 198, 8.11), (16, 201, 9.47), (16, 184, 8.71), (16, 186, 9.51), (16, 161, 8.35), (17, 162, 7.82), (17, 163, 6.17), (17, 164, 4.78), (17, 165, 6.14), (17, 22, 6.81), (17, 210, 4.68), (17, 23, 6.78), (17, 182, 10.27), (17, 229, 10.12), (17, 249, 9.74), (17, 252, 10.3), (17, 152, 9.77), (17, 151, 10.98), (17, 155, 11.8), (18, 22, 5.69), (18, 209, 7.77), (18, 210, 6.08), (18, 165, 5.12), (18, 211, 5.18), (18, 212, 7.87), (18, 216, 7.73), (18, 23, 6.71), (18, 166, 6.1), (18, 194, 9.19), (19, 164, 6.31), (19, 165, 6.27), (19, 166, 5.55), (19, 23, 4.98), (19, 24, 6.09), (19, 212, 9.54), (19, 142, 9.36), (19, 176, 9.12), (19, 168, 8.7), (19, 172, 10.47), (19, 173, 8.52), (19, 122, 12.34), (20, 166, 7.44), (20, 24, 5.46), (20, 142, 7.59), (20, 212, 8.4), (20, 216, 10.57), (20, 26, 4.96), (20, 168, 9.15), (21, 26, 3.63), (21, 142, 8.68), (21, 212, 6.33), (21, 145, 8.88), (21, 229, 8.82), (21, 247, 9.17), (22, 145, 8.08), (22, 229, 6.83), (22, 148, 8.5), (22, 249, 7.83), (22, 210, 7.31), (22, 211, 5.69), (22, 212, 5.95), (22, 216, 10.48), (23, 148, 5.6), (23, 249, 7.66), (23, 144, 5.74), (23, 164, 8.4), (23, 176, 9.39), (23, 180, 9.64), (23, 143, 8.52), (23, 145, 6.02), (23, 152, 9.58), (23, 163, 10.59), (23, 182, 11.72), (24, 145, 5.11), (24, 143, 5.29), (24, 144, 4.01), (24, 142, 5.98), (24, 176, 9.73), (25, 142, 6.37), (25, 143, 5.91), (25, 145, 5.6), (25, 247, 9.09), (25, 29, 9.15), (25, 144, 6.11), (25, 146, 9.11), (26, 142, 6.15), (27, 142, 5.51), (27, 141, 8.12), (27, 143, 7.6), (27, 64, 11.07), (28, 141, 5.01), (28, 57, 6.48), (28, 58, 6.57), (28, 61, 5.71), (28, 140, 6.01), (28, 64, 10.06), (28, 142, 4.26), (29, 140, 5.39), (29, 141, 5.58), (29, 143, 6.86), (29, 61, 5.0), (29, 139, 8.43), (29, 64, 8.31), (29, 68, 11.4), (30, 61, 5.91), (30, 65, 6.5), (30, 139, 5.84), (30, 140, 4.47), (30, 44, 9.4), (30, 138, 7.04), (30, 58, 8.26), (30, 62, 6.11), (31, 138, 7.8), (31, 139, 5.61), (31, 140, 5.59), (31, 141, 6.51), (31, 143, 8.22), (31, 123, 8.17), (32, 138, 5.76), (32, 139, 4.86), (32, 118, 8.4), (32, 137, 7.9), (32, 117,

11.27), (32, 121, 10.92), (32, 123, 8.99), (33, 43, 7.48), (33, 137, 6.19), (33, 138, 4.93), (33, 40, 5.76), (33, 44, 7.4), (33, 65, 10.71), (33, 69, 12.91), (33, 139, 6.61), (34, 43, 8.2), (34, 137, 4.98), (34, 138, 6.28), (34, 135, 8.73), (34, 136, 5.34), (34, 114, 8.3), (34, 118, 10.21), (35, 43, 6.17), (35, 135, 5.95), (35, 136, 4.09), (35, 39, 5.75), (35, 40, 5.64), (36, 135, 5.65), (36, 136, 5.17), (36, 134, 7.67), (36, 108, 10.69), (36, 111, 8.6), (36, 110, 9.63), (37, 135, 7.1), (37, 42, 9.51), (37, 130, 12.55), (37, 43, 8.99), (39, 43, 6.33), (40, 44, 6.09), (40, 71, 11.37), (41, 71, 7.77), (41, 45, 6.05), (41, 69, 8.29), (41, 70, 9.98), (41, 72, 10.64), (42, 46, 6.09), (42, 47, 8.57), (42, 130, 11.0), (42, 135, 8.93), (43, 128, 7.29), (43, 135, 6.5), (43, 47, 6.33), (43, 48, 8.64), (43, 138, 8.19), (43, 136, 6.51), (43, 137, 8.18), (44, 62, 7.04), (44, 138, 7.8), (44, 71, 8.43), (44, 48, 6.0), (44, 66, 6.19), (44, 73, 10.04), (44, 61, 10.7), (44, 65, 7.88), (44, 69, 9.37), (45, 71, 6.45), (45, 73, 6.72), (45, 49, 6.22), (45, 92, 10.71), (45, 72, 7.65), (45, 74, 9.0), (46, 92, 6.91), (46, 128, 7.64), (46, 135, 8.91), (46, 50, 6.51), (46, 51, 8.74), (46, 94, 8.77), (46, 130, 7.91), (47, 128, 6.3), (47, 94, 6.19), (47, 51, 6.06), (47, 62, 6.92), (47, 138, 8.36), (47, 63, 7.91), (47, 87, 9.32), (47, 92, 7.56), (48, 62, 5.59), (48, 63, 4.71), (48, 73, 7.0), (48, 66, 6.4), (48, 71, 8.99), (48, 52, 5.78), (48, 74, 9.07), (49, 73, 5.72), (49, 74, 6.13), (49, 76, 6.36), (49, 92, 8.36), (49, 79, 9.37), (49, 89, 7.83), (49, 72, 9.19), (50, 92, 6.28), (50, 76, 6.76), (50, 79, 6.73), (50, 89, 5.08), (50, 87, 6.55), (50, 94, 6.87), (50, 54, 8.22), (50, 88, 6.44), (51, 94, 7.49), (51, 87, 6.67), (51, 79, 8.1), (51, 58, 8.39), (51, 59, 4.93), (51, 62, 7.34), (51, 63, 6.82), (51, 86, 8.89), (51, 126, 11.52), (51, 56, 8.37), (52, 76, 7.84), (52, 79, 8.45), (52, 63, 6.44), (52, 73, 8.31), (52, 74, 8.98), (52, 75, 7.59), (52, 67, 11.12), (53, 76, 8.55), (53, 79, 6.84), (53, 81, 8.93), (53, 75, 8.98), (53, 77, 10.58), (53, 78, 9.27), (54, 81, 7.26), (54, 79, 8.29), (54, 87, 7.73), (54, 59, 5.81), (54, 60, 7.6), (55, 60, 6.76), (56, 60, 6.35), (57, 61, 6.22), (57, 140, 9.23), (58, 140, 6.45), (58, 62, 6.25), (58, 63, 8.62), (58, 124, 8.33), (58, 94, 10.21), (58, 126, 9.0), (59, 63, 6.05), (59, 64, 8.35), (60, 64, 5.84), (61, 140, 7.13), (61, 65, 6.11), (62, 140, 7.92), (62, 66, 6.28), (62, 138, 8.27), (62, 94, 10.44), (62, 126, 10.5), (63, 73, 8.76), (63, 67, 6.3), (64, 68, 6.22), (65, 69, 6.42), (65, 71, 8.87), (66, 71, 5.3), (66, 73, 8.08), (66, 70, 6.44), (67, 73, 8.36), (67, 71, 5.73), (75, 90, 10.2), (76, 90, 6.59), (76, 89, 6.28), (77, 90, 6.9), (78, 90, 6.42), (78, 88, 8.18), (78, 89, 5.68), (79, 88, 5.68), (79, 89, 4.55), (79, 87, 6.87), (80, 87, 5.74), (80, 88, 5.15), (80, 86, 8.68), (81, 86, 6.08), (81, 87, 4.58), (82, 86, 5.35), (82, 87, 5.95), (82, 99, 7.99), (82, 88, 7.78), (82, 102, 10.57), (82, 103, 11.84), (83, 98, 10.95), (84, 96, 8.31), (84, 98, 8.89), (85, 96, 5.52), (85, 99, 5.19), (85, 94, 8.56), (85, 95, 5.5), (85, 98, 6.12), (85, 102, 9.49), (86, 99, 6.46), (86, 94, 5.64), (86, 95, 3.97), (87, 94, 5.3), (87, 99, 8.23), (87, 92, 8.31), (87, 93, 5.9), (88, 92, 5.9), (88, 93, 4.66), (88, 103, 9.72), (88, 99, 8.77), (90, 131, 8.69), (91, 131, 4.97), (91, 129, 8.08), (91, 130, 5.61), (91, 132, 7.45), (92, 129, 5.66), (92, 130, 4.52), (92, 128, 7.28), (93, 128, 5.7), (93, 129, 5.01), (93, 99, 7.7), (93, 100, 7.69), (93, 105, 9.38), (93, 127, 7.74), (93, 103, 8.81), (93, 131, 8.51), (93, 132, 8.45), (94, 99, 7.94), (94, 127, 5.7), (94, 128, 4.65), (94, 126, 6.33), (95, 100, 7.35), (95, 126, 4.9), (95, 127, 5.42), (95, 99, 5.82), (95, 125, 8.36), (96, 125, 7.63), (96, 127, 6.02), (96, 100, 6.73), (97, 115, 9.7), (97, 127, 7.33), (97, 107, 7.69), (97, 101, 6.15), (97, 112, 9.0), (98, 102, 6.1), (99, 103, 6.49), (99, 105, 8.48), (99, 127, 8.95), (100, 105, 4.88), (100, 107, 7.42), (100, 104, 6.13), (100, 127, 7.89), (100, 134, 8.78), (100, 129, 8.01), (101, 105, 5.94), (103, 132, 7.46), (104, 132, 7.18), (105, 132, 5.15), (105, 127, 10.2), (105, 129, 7.65), (105, 134, 7.15), (105, 133, 6.05), (106, 132, 7.81), (106, 134, 7.42), (106, 133, 7.02), (107, 111, 6.28), (107, 134, 7.02), (107, 112, 6.91), (107, 115, 10.48), (107, 127, 8.84), (108, 134, 8.74), (108, 112, 6.32), (109, 113, 6.14), (110, 114, 6.25), (110, 137, 10.95), (111, 137, 7.57), (111, 115, 6.18), (111, 127, 8.41), (111, 134, 8.62), (111, 136, 7.76), (112, 116, 6.23), (113, 117, 6.1), (113, 120, 10.5), (114, 118, 6.33), (114, 137, 7.99), (114, 119, 8.66), (114, 125, 8.99), (115, 137, 7.4), (115, 125, 6.01), (115, 119, 6.1), (115, 120, 8.58), (115, 123, 9.1), (115, 127, 8.37), (115, 126,

7.93), (116, 120, 6.1), (117, 121, 6.56), (118, 123, 5.45), (118, 125, 6.37), (118, 139, 6.84), (118, 137, 9.46), (119, 125, 6.52), (119, 123, 4.98), (119, 171, 7.89), (119, 175, 9.5), (119, 124, 6.13), (120, 171, 6.49), (120, 175, 6.99), (120, 178, 11.44), (121, 175, 5.73), (121, 141, 9.18), (122, 175, 5.81), (122, 171, 6.38), (122, 172, 3.99), (122, 176, 7.19), (122, 140, 9.21), (122, 141, 7.08), (122, 142, 8.9), (123, 140, 5.68), (123, 141, 4.85), (123, 139, 5.85), (124, 139, 5.56), (124, 140, 5.17), (124, 141, 6.07), (124, 138, 8.32), (125, 138, 5.66), (125, 139, 4.26), (125, 137, 6.53), (126, 137, 5.62), (126, 138, 5.34), (126, 140, 8.12), (126, 136, 8.65), (127, 136, 6.06), (127, 137, 4.58), (127, 134, 7.01), (127, 135, 7.9), (128, 134, 5.89), (128, 136, 5.38), (128, 137, 5.84), (128, 138, 7.26), (128, 135, 5.66), (128, 133, 8.57), (129, 133, 5.21), (129, 134, 4.06), (129, 135, 5.14), (130, 135, 6.49), (142, 176, 10.42), (144, 148, 6.39), (144, 149, 8.06), (144, 179, 9.52), (144, 176, 8.37), (144, 180, 9.25), (145, 247, 8.85), (145, 249, 8.17), (145, 149, 7.8), (145, 150, 9.79), (145, 229, 12.27), (146, 150, 7.84), (147, 250, 6.1), (147, 151, 5.98), (147, 248, 5.77), (147, 249, 5.42), (147, 247, 8.7), (148, 249, 5.87), (148, 180, 7.74), (148, 152, 6.17), (148, 153, 8.63), (149, 180, 5.71), (149, 153, 6.17), (149, 179, 8.2), (149, 181, 8.71), (150, 250, 7.21), (150, 154, 6.12), (150, 253, 9.57), (151, 250, 4.92), (151, 253, 6.35), (151, 155, 6.2), (151, 249, 5.75), (151, 252, 7.54), (151, 162, 9.75), (152, 162, 6.58), (152, 180, 6.79), (152, 156, 6.21), (152, 157, 8.59), (152, 160, 9.27), (152, 181, 7.97), (152, 182, 7.68), (153, 157, 6.18), (153, 180, 7.93), (153, 181, 8.19), (154, 253, 7.03), (154, 158, 6.23), (154, 257, 10.13), (155, 253, 6.39), (155, 160, 5.44), (155, 162, 6.99), (155, 159, 6.73), (155, 252, 8.8), (155, 256, 8.9), (156, 162, 6.66), (156, 160, 5.09), (156, 161, 6.13), (156, 183, 8.81), (156, 180, 10.63), (156, 181, 9.29), (156, 182, 8.8), (158, 260, 10.42), (158, 257, 8.72), (158, 253, 9.04), (158, 256, 9.41), (160, 260, 12.62), (160, 183, 9.35), (160, 256, 10.21), (161, 183, 5.82), (161, 184, 7.89), (161, 205, 8.97), (162, 183, 5.42), (162, 184, 6.77), (162, 181, 8.54), (162, 182, 5.48), (163, 182, 5.45), (163, 184, 4.63), (164, 182, 7.34), (164, 184, 6.68), (164, 185, 4.98), (164, 186, 6.2), (164, 176, 8.91), (164, 177, 7.45), (164, 180, 9.75), (164, 172, 11.54), (164, 173, 7.97), (165, 186, 4.73), (165, 189, 7.2), (165, 194, 7.45), (165, 209, 9.52), (165, 216, 10.3), (165, 193, 10.31), (165, 197, 8.51), (165, 198, 9.17), (166, 186, 6.39), (166, 187, 4.95), (166, 189, 5.75), (166, 194, 8.41), (166, 188, 6.18), (166, 173, 7.17), (166, 185, 8.28), (167, 194, 9.14), (167, 188, 4.26), (167, 189, 4.45), (168, 187, 5.87), (168, 172, 6.41), (168, 173, 6.44), (168, 188, 5.22), (169, 188, 5.28), (169, 173, 6.32), (169, 174, 8.72), (170, 187, 5.41), (170, 188, 6.76), (170, 174, 5.87), (170, 185, 8.72), (170, 186, 8.15), (171, 175, 6.31), (172, 176, 6.59), (172, 177, 8.71), (173, 185, 6.92), (173, 186, 7.75), (173, 187, 6.06), (173, 177, 6.1), (173, 182, 10.36), (174, 185, 7.14), (174, 178, 6.15), (175, 179, 6.3), (176, 180, 6.13), (176, 181, 8.22), (176, 182, 8.42), (177, 182, 5.05), (177, 185, 6.6), (177, 181, 5.62), (177, 184, 7.7), (184, 201, 10.43), (186, 197, 8.09), (186, 201, 10.81), (188, 193, 9.81), (189, 193, 6.09), (189, 194, 6.39), (189, 197, 8.45), (190, 194, 6.71), (191, 215, 5.54), (191, 195, 8.31), (191, 218, 8.55), (191, 219, 8.71), (192, 218, 7.36), (192, 196, 7.92), (193, 197, 6.41), (194, 219, 5.56), (194, 198, 6.28), (194, 209, 9.93), (194, 215, 8.75), (194, 216, 7.4), (195, 219, 4.15), (195, 222, 5.81), (195, 218, 5.73), (195, 199, 6.1), (195, 202, 10.4), (196, 200, 6.26), (197, 201, 6.47), (198, 207, 7.18), (198, 219, 6.53), (198, 222, 6.81), (198, 202, 6.29), (198, 209, 7.72), (198, 223, 6.33), (199, 222, 6.83), (199, 203, 6.89), (200, 204, 8.1), (201, 205, 5.83), (201, 207, 7.02), (202, 207, 6.06), (202, 224, 7.99), (202, 206, 5.84), (202, 222, 9.22), (202, 223, 7.78), (205, 224, 9.35), (206, 224, 6.12), (207, 224, 5.34), (207, 223, 5.56), (207, 225, 6.49), (208, 223, 5.73), (208, 225, 4.48), (208, 227, 7.03), (208, 256, 9.79), (209, 223, 5.69), (209, 225, 6.29), (209, 226, 5.13), (209, 227, 5.95), (209, 219, 6.85), (209, 220, 5.62), (209, 216, 8.8), (210, 220, 7.07), (210, 227, 4.76), (210, 216, 8.36), (210, 229, 6.89), (210, 244, 10.01), (210, 249, 9.14), (210, 252, 8.18), (211, 220, 7.25), (211, 227, 6.58), (211, 228, 5.26), (211, 229, 5.92), (211, 216, 5.81), (211,

236, 7.58), (212, 216, 6.96), (212, 228, 5.49), (212, 229, 5.04), (212, 232, 4.41), (212, 233, 6.83), (212, 236, 6.27), (213, 233, 5.73), (213, 217, 7.08), (213, 220, 9.95), (213, 236, 5.79), (214, 233, 5.88), (214, 235, 5.68), (214, 236, 5.34), (214, 218, 8.5), (215, 219, 8.21), (216, 220, 6.0), (216, 236, 6.71), (216, 239, 8.83), (217, 236, 5.06), (217, 235, 6.66), (217, 239, 5.55), (217, 221, 6.17), (218, 222, 6.19), (219, 223, 6.19), (219, 226, 7.72), (220, 225, 7.19), (220, 226, 4.2), (220, 236, 7.65), (220, 239, 6.18), (220, 240, 5.72), (220, 241, 8.19), (220, 228, 7.61), (220, 235, 10.44), (220, 227, 6.72), (221, 239, 5.78), (223, 241, 9.59), (224, 241, 9.15), (225, 241, 6.21), (225, 242, 7.66), (225, 256, 10.31), (226, 241, 5.11), (226, 239, 7.17), (226, 240, 4.39), (226, 242, 6.32), (227, 242, 4.73), (227, 244, 7.32), (227, 252, 7.57), (227, 255, 9.43), (227, 256, 10.59), (228, 240, 5.28), (228, 242, 6.22), (228, 243, 4.74), (228, 244, 6.12), (228, 236, 6.01), (228, 237, 5.71), (229, 244, 4.96), (229, 246, 7.04), (229, 247, 6.05), (229, 248, 7.69), (229, 249, 7.59), (229, 252, 8.4), (230, 243, 7.52), (230, 244, 5.56), (230, 246, 4.89), (230, 245, 5.06), (230, 247, 4.37), (230, 237, 8.79), (231, 245, 6.62), (232, 237, 6.16), (232, 236, 5.9), (233, 237, 6.37), (233, 238, 8.63), (234, 238, 6.1), (235, 239, 6.21), (236, 240, 6.25), (236, 243, 8.72), (237, 243, 6.04), (242, 255, 8.24), (244, 248, 8.04), (244, 251, 6.65), (244, 249, 8.7), (244, 252, 7.61), (244, 255, 8.85), (245, 251, 8.74), (246, 251, 7.51), (248, 252, 7.98), (249, 253, 7.24), (249, 254, 9.02), (250, 254, 6.25), (251, 255, 6.04), (252, 256, 6.5), (252, 257, 8.69), (253, 257, 6.06), (253, 258, 8.47), (254, 258, 5.9), (255, 259, 6.24), (256, 260, 6.31), (257, 261, 6.16), (258, 262, 6.11), (259, 263, 6.21), (260, 264, 6.19), (261, 265, 6.88)]

Qclosed, N=838 (Residue i, Residue j, σij )

[(0, 242, 4.83), (0, 243, 6.67), (0, 5, 6.77), (0, 241, 6.29), (0, 6, 8.27), (1, 5, 7.4), (1, 241, 5.06), (1, 243, 7.66), (1, 240, 7.43), (1, 238, 10.45), (1, 239, 9.14), (1, 235, 10.58), (1, 236, 11.54), (2, 241, 5.7), (3, 7, 8.32), (4, 8, 6.17), (4, 9, 8.53), (4, 225, 9.89), (5, 225, 7.15), (5, 9, 5.99), (5, 242, 7.07), (5, 259, 8.34), (5, 241, 6.1), (6, 259, 4.71), (6, 262, 7.72), (6, 10, 6.29), (6, 263, 7.8), (6, 258, 7.72), (6, 242, 9.05), (6, 255, 7.96), (7, 263, 6.9), (7, 11, 6.92), (8, 12, 6.61), (8, 206, 8.58), (8, 225, 6.66), (8, 224, 7.2), (9, 225, 6.63), (9, 160, 11.05), (9, 256, 5.7), (9, 260, 7.96), (9, 242, 9.49), (9, 255, 7.86), (9, 259, 5.98), (10, 259, 5.34), (10, 260, 5.36), (10, 256, 6.33), (10, 263, 7.33), (11, 206, 6.66), (11, 263, 10.09), (12, 206, 4.87), (12, 160, 7.33), (12, 260, 10.4), (12, 256, 8.42), (12, 207, 5.78), (12, 208, 6.28), (12, 224, 7.87), (12, 225, 7.74), (13, 206, 5.17), (13, 208, 7.97), (13, 159, 7.69), (13, 160, 5.08), (13, 205, 5.68), (13, 161, 6.14), (13, 204, 9.08), (14, 205, 6.18), (14, 206, 6.24), (14, 207, 5.02), (14, 160, 5.44), (14, 161, 4.43), (14, 208, 6.17), (14, 184, 9.61), (14, 201, 8.63), (15, 160, 7.02), (15, 161, 6.21), (15, 162, 4.97), (15, 208, 4.69), (15, 155, 8.11), (15, 163, 6.18), (15, 210, 7.14), (15, 227, 9.62), (15, 252, 9.37), (15, 256, 10.23), (16, 207, 7.28), (16, 208, 6.13), (16, 209, 4.88), (16, 210, 5.96), (16, 163, 4.51), (16, 198, 8.43), (16, 201, 9.79), (16, 165, 6.61), (16, 184, 8.68), (16, 186, 9.29), (16, 161, 8.38), (17, 162, 7.82), (17, 163, 6.14), (17, 164, 4.71), (17, 165, 6.11), (17, 22, 6.47), (17, 210, 4.67), (17, 23, 6.91), (17, 229, 9.81), (17, 249, 9.61), (17, 252, 10.14), (17, 152, 9.66), (17, 151, 10.78), (17, 155, 11.82), (18, 22, 5.53), (18, 209, 7.55), (18, 210, 6.03), (18, 165, 5.0), (18, 23, 6.78), (18, 211, 5.13), (18, 216, 8.31), (18, 166, 6.0), (18, 194, 9.25), (19, 164, 6.23), (19, 165, 6.07), (19, 166, 5.54), (19, 23, 4.9), (19, 24, 6.39), (19, 144, 8.89), (19, 168, 8.62), (19, 176, 9.65), (19, 122, 12.15), (19, 124, 12.28), (19, 172, 10.93), (20, 166, 7.32), (20, 24, 5.77), (20, 212, 8.56), (20, 57, 8.65), (21, 211, 5.05), (21, 212, 4.76), (21, 216, 7.82), (21, 145, 11.44), (21, 210, 8.15), (21, 229, 7.84), (22, 229, 6.31), (22, 145, 8.54), (22, 249, 7.83), (22, 210, 6.99), (22, 211, 5.53), (22, 216, 10.55), (22,

227, 10.15), (23, 145, 5.86), (23, 148, 6.16), (23, 249, 7.78), (23, 143, 8.64), (23, 144, 5.79), (23, 180, 10.62), (23, 163, 11.01), (23, 164, 8.56), (23, 176, 10.18), (23, 182, 12.53), (23, 152, 10.41), (24, 143, 5.56), (24, 144, 4.28), (24, 145, 5.19), (24, 142, 6.2), (24, 176, 10.83), (25, 142, 6.63), (25, 143, 5.83), (25, 145, 5.27), (25, 144, 5.97), (25, 146, 8.63), (26, 142, 6.67), (26, 60, 10.11), (27, 142, 5.6), (27, 60, 8.94), (27, 141, 8.41), (27, 143, 7.18), (28, 141, 5.42), (28, 142, 4.18), (28, 57, 5.91), (28, 60, 7.28), (28, 61, 5.33), (28, 64, 9.5), (28, 140, 6.44), (29, 140, 5.38), (29, 141, 5.63), (29, 61, 4.45), (29, 64, 7.81), (29, 65, 7.31), (29, 139, 8.46), (30, 61, 5.67), (30, 65, 6.51), (30, 139, 5.77), (30, 140, 4.38), (30, 44, 9.73), (30, 138, 7.14), (30, 58, 8.24), (30, 62, 6.16), (31, 139, 5.62), (31, 140, 5.51), (31, 141, 6.46), (31, 123, 8.23), (32, 139, 4.95), (32, 137, 8.16), (32, 138, 5.83), (32, 118, 8.33), (32, 117, 11.44), (32, 121, 11.03), (32, 123, 9.38), (33, 43, 7.66), (33, 137, 6.36), (33, 138, 4.84), (33, 40, 6.07), (33, 44, 7.4), (33, 65, 10.31), (33, 69, 12.57), (33, 139, 6.58), (34, 43, 8.16), (34, 137, 5.21), (34, 138, 6.21), (34, 135, 8.89), (34, 136, 5.57), (34, 114, 8.9), (34, 118, 10.38), (35, 43, 6.12), (35, 135, 5.98), (35, 136, 4.06), (35, 39, 5.81), (35, 40, 5.61), (36, 135, 5.71), (36, 136, 5.05), (36, 134, 7.8), (36, 110, 10.35), (36, 111, 8.95), (37, 135, 7.07), (37, 134, 9.97), (39, 43, 6.41), (40, 44, 6.17), (40, 71, 11.26), (41, 71, 7.66), (41, 45, 6.07), (41, 69, 8.1), (41, 72, 10.47), (41, 70, 9.77), (42, 46, 6.15), (42, 130, 10.9), (42, 135, 9.13), (43, 128, 7.35), (43, 135, 6.6), (43, 47, 6.35), (43, 48, 8.64), (43, 138, 8.32), (43, 136, 6.65), (43, 137, 8.22), (44, 62, 7.11), (44, 138, 8.0), (44, 48, 6.08), (44, 66, 6.2), (44, 71, 8.25), (44, 65, 7.93), (44, 69, 9.33), (45, 71, 6.34), (45, 49, 6.39), (45, 73, 6.47), (45, 74, 8.33), (45, 72, 7.54), (45, 130, 11.52), (46, 92, 6.99), (46, 50, 6.68), (46, 51, 8.73), (46, 94, 8.82), (46, 128, 7.82), (46, 130, 8.01), (46, 135, 9.13), (46, 91, 9.83), (46, 129, 8.7), (47, 128, 6.36), (47, 94, 6.16), (47, 62, 6.98), (47, 51, 6.0), (47, 63, 8.11), (47, 87, 9.25), (47, 92, 7.51), (47, 138, 8.56), (48, 62, 5.71), (48, 63, 4.99), (48, 52, 5.55), (48, 73, 7.16), (48, 66, 6.53), (48, 71, 8.92), (49, 73, 5.89), (49, 53, 8.26), (49, 74, 6.23), (49, 77, 7.3), (49, 79, 8.99), (49, 89, 8.02), (49, 92, 8.32), (49, 72, 9.39), (49, 75, 8.1), (50, 92, 6.34), (50, 79, 6.41), (50, 89, 5.48), (50, 54, 8.31), (50, 87, 6.31), (50, 88, 6.37), (50, 94, 6.92), (51, 94, 7.52), (51, 87, 6.62), (51, 59, 4.85), (51, 63, 6.89), (51, 58, 8.11), (51, 62, 7.29), (51, 56, 8.43), (51, 126, 11.74), (52, 79, 8.6), (52, 63, 6.31), (52, 73, 8.66), (52, 76, 7.87), (53, 79, 7.22), (53, 77, 8.78), (53, 76, 8.36), (54, 81, 7.83), (54, 59, 5.77), (54, 60, 7.5), (54, 87, 8.03), (55, 60, 6.55), (56, 60, 6.2), (57, 61, 6.1), (57, 140, 8.61), (58, 140, 6.33), (58, 62, 6.27), (58, 124, 8.11), (58, 126, 8.65), (58, 94, 9.73), (59, 63, 6.03), (59, 64, 8.37), (60, 64, 5.99), (61, 65, 6.16), (61, 140, 6.82), (62, 140, 7.91), (62, 66, 6.24), (62, 138, 8.34), (62, 94, 10.45), (62, 126, 10.69), (63, 67, 6.24), (63, 73, 9.4), (64, 68, 6.35), (65, 69, 6.57), (65, 71, 8.85), (66, 71, 5.24), (66, 73, 8.44), (66, 70, 6.28), (77, 89, 6.49), (77, 90, 7.37), (77, 92, 10.13), (78, 89, 5.94), (78, 90, 7.35), (78, 88, 8.38), (79, 88, 5.45), (79, 89, 4.63), (79, 87, 6.69), (80, 88, 5.1), (80, 86, 8.7), (80, 87, 5.82), (80, 103, 13.16), (81, 86, 5.98), (81, 87, 4.57), (82, 86, 5.38), (82, 87, 6.16), (82, 88, 7.88), (82, 99, 7.99), (82, 102, 10.16), (82, 103, 11.75), (83, 98, 10.48), (83, 191, 6.39), (83, 192, 9.57), (84, 98, 8.29), (84, 191, 6.03), (84, 96, 8.25), (84, 167, 6.96), (85, 96, 5.59), (85, 98, 5.89), (85, 99, 4.94), (85, 94, 8.48), (85, 95, 5.51), (85, 102, 9.1), (85, 103, 11.18), (86, 99, 6.32), (86, 94, 5.69), (86, 95, 4.2), (87, 94, 5.42), (87, 99, 8.33), (87, 92, 8.2), (87, 93, 5.98), (88, 92, 5.91), (88, 93, 4.91), (88, 103, 9.86), (88, 99, 9.01), (89, 93, 6.55), (90, 131, 8.39), (91, 131, 4.65), (91, 129, 7.92), (91, 130, 5.52), (91, 132, 7.08), (92, 129, 5.87), (92, 130, 4.92), (92, 128, 7.4), (93, 128, 5.69), (93, 129, 5.02), (93, 99, 7.79), (93, 100, 7.73), (93, 127, 7.84), (93, 103, 8.88), (93, 105, 9.47), (94, 99, 7.89), (94, 127, 5.76), (94, 128, 4.61), (94, 126, 6.56), (94, 138, 9.09), (95, 100, 7.32), (95, 127, 5.4), (95, 99, 5.81), (95, 126, 4.9), (95, 125, 8.4), (96, 125, 7.6), (96, 127, 5.95), (96, 100, 6.63), (96, 126, 5.21), (96, 167, 11.32), (97, 115, 10.0), (97, 127, 7.44), (97, 107, 7.59), (97, 101, 6.05), (97, 112, 8.93), (98, 102, 6.13), (98,

190, 11.01), (98, 191, 12.63), (99, 103, 6.64), (99, 105, 8.44), (99, 127, 8.95), (100, 105, 4.84), (100, 104, 6.11), (100, 107, 7.46), (100, 127, 7.91), (100, 134, 8.87), (100, 129, 8.02), (103, 132, 7.47), (104, 132, 7.43), (105, 132, 5.38), (105, 127, 10.24), (105, 129, 7.77), (105, 133, 6.09), (105, 134, 7.32), (106, 132, 8.01), (106, 134, 7.42), (106, 133, 6.96), (107, 111, 6.3), (107, 134, 7.16), (107, 112, 6.86), (107, 115, 10.42), (107, 127, 8.84), (108, 134, 9.08), (108, 112, 6.27), (109, 113, 6.05), (110, 114, 6.26), (111, 137, 7.82), (111, 115, 6.11), (111, 127, 8.61), (111, 134, 8.75), (112, 116, 6.18), (112, 127, 9.59), (112, 119, 10.4), (112, 171, 12.59), (113, 117, 6.2), (114, 137, 8.34), (114, 118, 6.49), (114, 119, 8.76), (114, 125, 9.15), (115, 137, 7.38), (115, 125, 5.93), (115, 119, 6.2), (115, 126, 7.78), (115, 127, 8.39), (116, 171, 7.28), (116, 120, 6.16), (117, 121, 6.83), (118, 123, 5.87), (118, 139, 6.89), (118, 125, 6.45), (118, 137, 9.67), (119, 125, 6.22), (119, 123, 4.7), (119, 172, 5.08), (119, 171, 5.54), (119, 169, 7.26), (119, 175, 8.67), (119, 124, 5.91), (119, 170, 8.23), (119, 188, 12.6), (120, 171, 5.22), (120, 175, 6.42), (120, 178, 11.39), (121, 141, 9.25), (122, 171, 7.16), (122, 172, 4.05), (122, 175, 5.82), (122, 140, 9.37), (122, 141, 7.25), (122, 142, 9.08), (122, 176, 6.77), (123, 169, 8.6), (123, 140, 5.81), (123, 141, 4.86), (123, 168, 9.29), (123, 172, 5.83), (123, 139, 6.22), (124, 139, 5.52), (124, 140, 5.13), (124, 141, 6.01), (124, 168, 7.03), (124, 169, 6.98), (124, 138, 8.36), (124, 172, 7.17), (125, 169, 8.57), (125, 138, 5.72), (125, 139, 4.24), (125, 137, 6.68), (126, 137, 5.67), (126, 138, 5.43), (126, 140, 8.14), (126, 136, 8.75), (127, 136, 6.11), (127, 137, 4.53), (127, 134, 7.0), (127, 135, 7.84), (128, 134, 5.97), (128, 136, 5.49), (128, 137, 5.8), (128, 138, 7.29), (128, 133, 8.61), (128, 135, 5.67), (129, 133, 5.21), (129, 134, 4.12), (129, 135, 5.12), (130, 135, 6.2), (144, 148, 6.66), (144, 180, 9.76), (144, 176, 8.8), (144, 179, 10.29), (144, 164, 12.15), (145, 149, 8.0), (145, 150, 10.07), (145, 249, 8.11), (145, 247, 8.62), (145, 229, 12.27), (145, 246, 12.16), (146, 150, 8.05), (147, 250, 6.27), (147, 151, 6.02), (147, 248, 5.86), (147, 249, 5.54), (147, 247, 8.75), (148, 249, 5.97), (148, 180, 7.72), (148, 152, 6.18), (148, 153, 8.61), (149, 180, 5.73), (149, 153, 6.14), (149, 179, 8.33), (149, 181, 8.61), (150, 154, 6.0), (150, 157, 10.24), (150, 253, 9.48), (150, 250, 7.18), (151, 250, 4.82), (151, 253, 6.26), (151, 155, 6.21), (151, 162, 9.6), (151, 249, 5.82), (151, 252, 7.54), (152, 162, 6.45), (152, 180, 6.88), (152, 156, 6.2), (152, 160, 9.18), (152, 181, 7.89), (152, 182, 7.55), (152, 183, 9.75), (153, 157, 6.15), (153, 180, 7.93), (153, 181, 8.0), (154, 253, 7.16), (154, 158, 6.21), (154, 257, 10.01), (155, 253, 6.43), (155, 160, 5.36), (155, 159, 6.62), (155, 161, 7.24), (155, 162, 7.01), (155, 252, 8.98), (155, 256, 8.75), (156, 162, 6.71), (156, 160, 5.02), (156, 161, 5.97), (156, 183, 8.61), (156, 180, 10.7), (156, 181, 9.2), (158, 260, 10.21), (158, 257, 8.84), (158, 253, 9.32), (158, 256, 9.57), (159, 260, 12.32), (160, 183, 9.21), (160, 260, 12.35), (160, 256, 10.11), (161, 183, 5.68), (161, 184, 7.89), (161, 205, 9.1), (162, 183, 5.31), (162, 184, 6.66), (162, 181, 8.51), (162, 182, 5.47), (163, 182, 5.42), (163, 184, 4.68), (164, 182, 7.4), (164, 184, 6.87), (164, 185, 4.98), (164, 186, 6.17), (164, 176, 8.74), (164, 180, 9.66), (164, 172, 11.52), (164, 173, 7.94), (164, 177, 7.58), (165, 186, 4.7), (165, 194, 7.62), (165, 189, 7.71), (165, 209, 9.22), (165, 193, 10.61), (165, 197, 8.68), (165, 198, 9.31), (166, 186, 6.43), (166, 187, 4.82), (166, 189, 6.09), (166, 194, 8.15), (166, 188, 6.33), (166, 173, 7.79), (166, 185, 8.55), (167, 194, 8.71), (167, 188, 4.36), (167, 189, 4.48), (168, 187, 5.95), (168, 173, 7.32), (168, 188, 5.21), (168, 172, 7.65), (169, 188, 5.36), (169, 173, 6.3), (169, 187, 6.04), (170, 188, 6.29), (170, 187, 5.3), (170, 174, 5.98), (170, 185, 8.7), (170, 186, 7.85), (171, 175, 6.25), (171, 188, 9.92), (172, 176, 6.29), (172, 177, 8.57), (173, 177, 6.05), (173, 182, 10.24), (173, 185, 6.81), (173, 186, 7.61), (173, 187, 6.31), (174, 185, 6.96), (174, 178, 6.22), (175, 179, 6.36), (176, 180, 6.03), (176, 181, 8.19), (176, 182, 8.2), (177, 182, 4.93), (177, 181, 5.67), (177, 185, 6.29), (177, 184, 7.27), (184, 201, 10.42), (186, 197, 7.99), (186, 201, 10.7), (189, 193, 6.13), (189, 197, 8.22), (189, 194, 6.51), (190, 194, 7.78), (192, 196, 8.31), (192, 218, 8.08), (193, 197, 6.49), (193, 198,

8.8), (194, 198, 6.24), (194, 209, 9.76), (194, 222, 9.05), (194, 219, 5.7), (194, 216, 8.15), (194, 215, 11.18), (195, 219, 4.4), (195, 222, 5.72), (195, 199, 6.04), (195, 202, 10.39), (195, 218, 6.14), (196, 200, 6.26), (197, 201, 6.5), (198, 207, 7.27), (198, 202, 6.27), (198, 219, 6.93), (198, 222, 6.9), (198, 209, 7.78), (198, 223, 6.33), (199, 222, 6.95), (199, 203, 7.06), (200, 204, 8.25), (201, 205, 5.84), (201, 207, 7.2), (202, 207, 6.19), (202, 224, 8.13), (202, 206, 5.88), (202, 222, 9.36), (202, 223, 7.85), (205, 224, 9.32), (206, 224, 6.12), (207, 224, 5.33), (207, 223, 5.57), (207, 225, 6.43), (208, 223, 5.77), (208, 225, 4.55), (208, 227, 7.01), (208, 256, 10.08), (209, 223, 5.76), (209, 225, 6.43), (209, 226, 5.17), (209, 227, 6.09), (209, 220, 5.69), (209, 219, 6.82), (209, 216, 9.4), (210, 220, 7.23), (210, 227, 4.71), (210, 229, 6.87), (210, 244, 10.08), (210, 249, 9.43), (210, 252, 8.22), (211, 220, 7.16), (211, 227, 6.39), (211, 228, 5.1), (211, 229, 6.08), (211, 216, 6.14), (212, 216, 6.55), (212, 229, 5.23), (212, 228, 4.89), (212, 231, 7.35), (212, 238, 7.56), (212, 239, 8.35), (213, 227, 7.56), (213, 220, 7.33), (213, 228, 4.74), (213, 238, 4.59), (213, 239, 4.62), (213, 217, 5.46), (213, 237, 7.82), (214, 237, 7.17), (214, 238, 5.74), (216, 220, 6.47), (216, 221, 8.89), (217, 239, 5.85), (217, 221, 6.17), (218, 222, 6.14), (219, 223, 6.19), (219, 226, 7.7), (220, 225, 7.06), (220, 226, 4.18), (220, 240, 5.76), (220, 241, 8.2), (220, 239, 7.29), (220, 228, 7.91), (220, 227, 6.81), (223, 241, 9.85), (224, 241, 9.23), (225, 241, 6.26), (225, 242, 7.85), (225, 256, 10.67), (226, 241, 5.22), (226, 240, 4.53), (226, 239, 7.88), (226, 242, 6.43), (227, 242, 4.82), (227, 252, 7.74), (227, 244, 7.4), (227, 255, 9.44), (227, 256, 10.57), (228, 239, 6.77), (228, 240, 5.8), (228, 242, 6.3), (228, 243, 4.62), (228, 244, 5.99), (228, 238, 6.89), (229, 244, 4.94), (229, 246, 6.73), (229, 247, 5.97), (229, 248, 8.01), (229, 249, 7.51), (229, 252, 7.98), (230, 244, 5.89), (230, 246, 4.64), (230, 245, 5.11), (230, 247, 4.21), (230, 238, 9.65), (231, 244, 6.26), (231, 245, 4.8), (231, 238, 6.87), (231, 235, 6.18), (231, 243, 6.92), (232, 244, 8.0), (232, 245, 4.69), (233, 245, 6.42), (234, 238, 8.78), (234, 245, 8.47), (235, 243, 7.7), (235, 245, 8.01), (238, 243, 7.12), (239, 243, 7.55), (242, 255, 8.3), (244, 248, 8.73), (244, 251, 6.96), (244, 249, 9.03), (244, 252, 7.59), (244, 255, 8.73), (245, 251, 8.84), (246, 251, 7.41), (248, 252, 7.99), (249, 253, 7.19), (250, 254, 6.18), (251, 255, 6.19), (252, 256, 6.54), (252, 257, 8.64), (253, 257, 5.96), (254, 258, 5.83), (255, 259, 6.29), (256, 260, 6.35), (256, 261, 8.64), (257, 261, 6.17), (258, 262, 6.09), (259, 263, 6.07), (260, 264, 6.13), (261, 265, 7.18)]
